# Low-dimensional and monotonic preparatory activity in mouse anterior lateral motor cortex

**DOI:** 10.1101/203414

**Authors:** Hidehiko K. Inagaki, Miho Inagaki, Sandro Romani, Karel Svoboda

**Affiliations:** Janelia Research Campus, HHMI, Ashburn VA 20147

## Abstract

Neurons in multiple brain regions fire trains of action potentials anticipating specific movements, but this ‘preparatory activity’ has rarely been compared across behavioral tasks in the same brain region. We compared preparatory activity in auditory and tactile delayed-response tasks, with directional licking as the output. The anterior lateral motor cortex (ALM) is necessary for motor planning in both tasks. Multiple features of ALM preparatory activity during the delay epoch were similar across tasks. First, majority of neurons showed direction-selective activity and spatially intermingled neurons were selective for either movement direction. Second, many cells showed mixed coding of sensory stimulus and licking direction, with a bias toward licking direction. Third, delay activity was largely monotonic and low-dimensional. Fourth, pairs of neurons with similar direction selectivity showed high spike-count correlations. Our study forms the foundation to analyze the neural circuits underlying preparatory activity in a genetically tractable model organism.

## Introduction

Short-term memory is the ability of the brain to maintain information without external cues over times of seconds. Motor planning is a short-term memory that links past sensory information to future movements. Neurons in frontal and parietal cortex and related brain regions show persistent or ramping changes in spike rate during different types of short-term memory ^1-13^. Neural correlates of motor planning, referred to here as ‘preparatory activity’, is an example of such a memory trace. Preparatory activity anticipates movements and has selectivity for specific movements (such as saccade location, or movement direction of the hand, wrist or tongue).

Preparatory activity has been studied extensively in non-human primates ^1-7,14-27^. Preparatory activity has been detected in the primary motor cortex ^5,14,15^, the premotor or supplemental motor cortex ^5,15,16,15^, frontal eye field (FEF) ^3,17,19^, parietal cortex ^6,20,27^, striatum ^15,26^, superior colliculus ^24,25^, motor-related thalamus ^22^, and cerebellum ^23^. Neurons within a brain region show activity patterns with diverse dynamics. A subset of neurons ramp during the delay epoch up to the movement onset ^3-6,16-21^. Some neurons show selectivity only before the go cue. Other neurons show selectivity during the delay epoch as well as during the movement. A third class of neurons become selective only after the go cue in the peri-movement epoch, consistent with activity that might be causally related to movement execution.

More recently motor planning has been studied in rodents ^9^, including head-restrained mice performing a delayed response task ^8^. An instruction informs the type of action to be performed. A go cue determines the timing of the action. The instruction and go cue are separated by a delay epoch. In mice performing a tactile delayed-response licking task ^8^ (Fig. 1a), a large proportion of neurons in anterior lateral motor cortex (ALM) exhibit preparatory activity that predicts directional licking ^8,28^. Optogenetic manipulation has provided multiple lines of evidence supporting the view that preparatory activity in ALM is causally related to motor planning ^8,29,30^ {Svoboda and Li, *in press*}.

**Figure 1.**
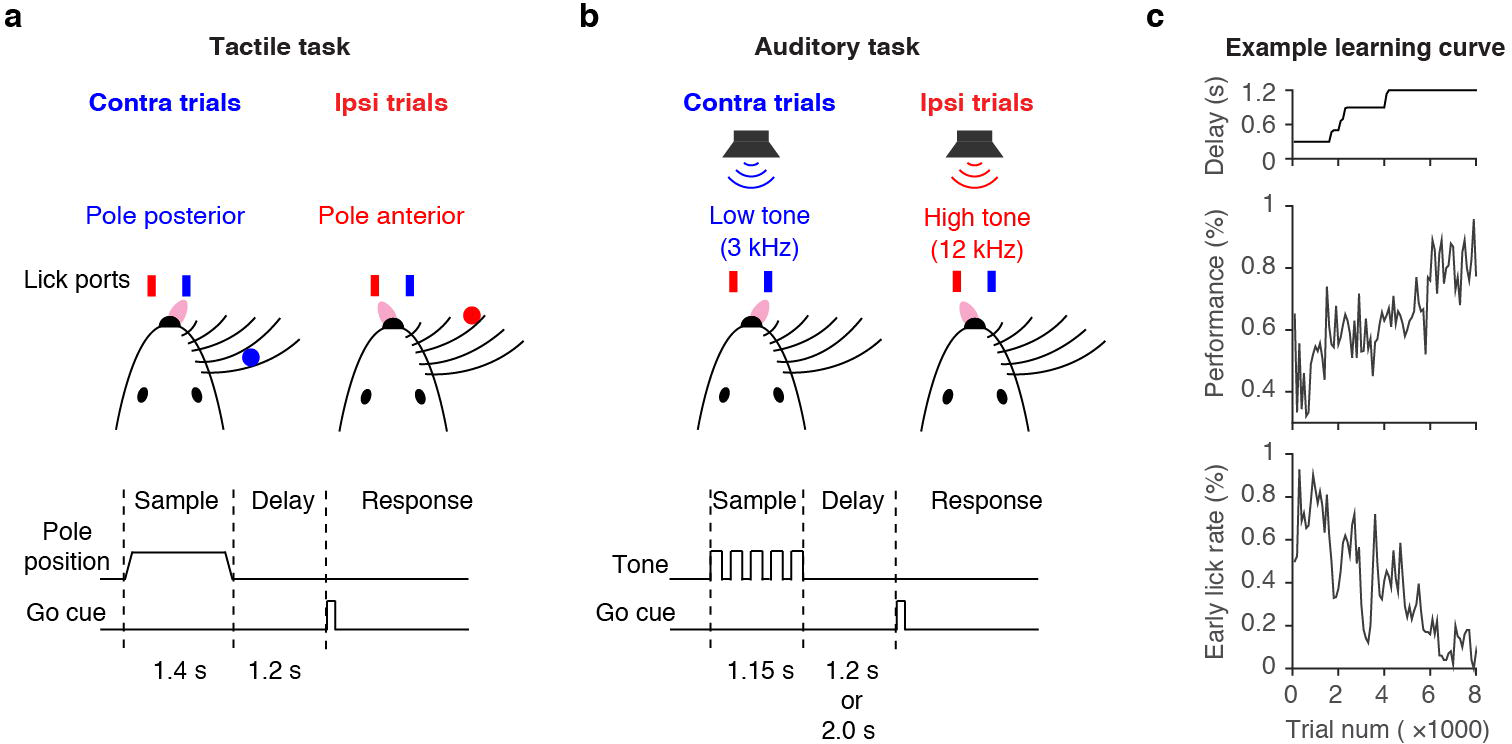
Behavioral tasks. **a.** Schematic, tactile task. During the delay epoch, an object is presented within reach of the whiskers in one of two locations. Mice report the location of the object by directional licking. **b.** Schematic, auditory task. Mice discriminate the frequency of tones presented during the sample epoch. **c.** Example learning curve, auditory task. The delay duration was gradually increased (top). Behavioral performance improved with training, while early lick rates decreased (middle and bottom).

Simultaneous recordings of multiple neurons have provided a comprehensive description of preparatory activity. Over time, populations of neurons trace out a trajectory in activity space, where each neuron corresponds to a dimension of activity space. Dimensionality reduction methods have revealed that these activity trajectories occupy a low-dimensional manifold, where the dimensionality scales with the dimensionality of the behavior ^31^. The detailed structure of the activity trajectories during motor planning predicts parameters of the future movements. Different endpoints of trajectories could reflect optimal ‘initial conditions’ for specific planned movements ^32^.

Here we explored the dynamics of populations of ALM neurons in several types of delayed response tasks in mice. The goal was to identify task-invariant features of the dynamics underlying motor planning. We developed an auditory delayed-response task and compared it to a previously reported tactile delayed-response task ^8^. We found that ALM is required for motor planning independent of the sensory modality. ALM neurons showed direction-selective ramping activity for both tasks. Many features of ALM activity were similar across the tasks, including direction selectivity, the properties of ramping activity, the dimensionality of the neural dynamics, and the spike count correlations among cells with similar and different selectivity. Some of these task-invariant features have also been reported in studies in behaving non-human and human primates ^3-5,33,34^. ALM thus provides a powerful model system to study the mechanism of motor planning in a genetically accessible organism.

## Results

### Task design

We developed an auditory delayed-response task for head-fixed mice and compared behavior and neural dynamics to the tactile delayed-response task (Fig. 1a vs. b). In the auditory task, the instruction was five tones played at one of two frequencies (3 or 12 kHz). In the tactile task, the instruction was a pole presented to either anterior or posterior location within reach of the whiskers. The instruction stimuli were presented during the sample epoch, which was followed by a delay epoch (1.2 or 2.0 seconds). The delay epoch was terminated by a brief (0.1 s) go cue (6 kHz for the sound task, 3.4 kHz for the tactile task). During the response epoch animals reported the instruction by licking either the right or left lick port. Premature licking responses resulted in a timeout (early-lick). Individual mice were trained for specific tasks ^35^. Criterion performance was 80 % correct trials with early-lick rate less than 20 % (Fig. 1c). Performance during recording was 86.1 ± 9.8 % (mean ± s.d.) with 7.0 ± 6.9 % (mean ± s.d.) early-lick rate (See Table 1 for detail). No mice were excluded based on behavioral performance.

**Table 1.** Summary of units and performance.

|  | Tactile task<br>( delay 1.2 s ) | Auditory task<br>( delay 1.2 s ) | Auditory task<br>( delay 2.0 s ) |
| --- | --- | --- | --- |
| # of animals | 7 | 6 | 6 |
| # of recording sessions | 30 | 37 | 20 |
| # of recording sessions for session based analysis* | 19 | 31 | 20 |
| # of units | 566 | 639 | 755 |
| # of putative pyramidal cells | 471 | 539 | 667 |
| # of selective pyramidal cells (Fig. 5a-c) | 431<br>91.5 % of pyramidal cells | 478<br>88.7 % of pyramidal cells | 618<br>92.7 % of pyramidal cells |
| # of delay-selective cells (Fig. 6a-c, top +middle) | 324<br>68.8 % of pyramidal cells | 340<br>63.1 % of pyramidal cells | 422<br>63.3 % of pyramidal cells |
| Performance** | 80.3 ± 10.1 %<br>(mean ± s.d.)<br>60.3 – 93.4 %<br>(min - max) | 91.4 ± 7.0 %<br>(mean ± s.d.)<br>69.2 – 99.7 %<br>(min - max) | 87.2 ± 8.4 %<br>(mean ± s.d.)<br>73.6 – 98.7 %<br>(min - max) |
| Early lick rate** | 5.7 ± 5.1 %<br>(mean ± s.d.) | 9.5 ± 9.4 %<br>(mean ± s.d.) | 5.4 ± 3.5 %<br>(mean ± s.d.) |
| # of trials ** | 357 ± 71 trials<br>(mean ± s.d.) | 343 ± 132 trials<br>(mean ± s.d.) | 307 ± 55 trials<br>(mean ± s.d.) |
| Depth | 397 – 985 um<br>(5 -95 % percentile) | 389 – 946 um<br>(5 -95 % percentile) | 365 – 984 um<br>(5 -95 % percentile) |

\*For the session based analysis (Fig. 15 and 16), recording sessions with more than 5 simultaneously recorded pyramidal cells were analyzed (see also supplementary file).
\*\* Behavioral performance, early lick rates and trial number were calculated based on behavioral session. In some cases (8 sessions in auditory tasks with 1.2 s delay), we moved probe within a behavioral session to record units from different depth. In this case we had two recording sessions per behavioral session.

We used logistic regression to test for correlations in behavioral performance across trials. We tested the effects of licking direction, performance (correct or incorrect), and optogenetic manipulation. Although there were significant effects of previous trials, the total variance explained by previous trials was less than 1 % in all cases (Table 2). Previous trials also had only minor effects on spike rates during the baseline period before the sample epoch (pre-sample epoch) (median of variance explained lower than 2.2 % in all tasks) (Table 3). We therefore treated trials as independent.

**Table 2.** Performance explained by previous trial types. Coefficients and variance explained of a logistic regression described below. logit(Outcome) ~ 1 + Prev_dir + Prev_correct + Stim Outcome, performance of current trial (correct 1, incorrect 0); Prev_dir, direction of previous trial; Prev_correct, performance of previous trials; Stim, optogentic manipulation in previous trial.

|  | Tactile task<br>( delay 1.2 s ) | Auditory task<br>( delay 1.2 s ) | Auditory task<br>( delay 2.0 s ) |
| --- | --- | --- | --- |
| Intercept | 1.38<br>$p = 2.47\text{e}^{-118}$ | 1.91<br>$p = 6.68\text{e}^{-88}$ | 1.68<br>$p = 1.03\text{e}^{-60}$ |
| Direction of previous trial<br>(same direction 1, not same 0) | -0.0763<br>$p = 0.145$ | 0.349<br>$p = 0.000132$ | -0.152<br>$p = 0.0652$ |
| Performance of previous trial<br>(correct 1, incorrect 0) | 0.194<br>$p = 0.00109$ | 0.465<br>$p = 4.61\text{e}^{-6}$ | 0.500<br>$p = 1.63\text{e}^{-7}$ |
| Optogenetic manipulation in<br>previous trial<br>(yes 1, no 0) | 0.0586<br>$p = 0.359$ | 0.216<br>$p = 0.070$ | -0.137<br>$p = 0.101$ |
| Variance explained | 0.14 % | 0.57 % | 0.67 % |
| Variance explained by<br>regression with interactions | 0.17 % | 0.59 % | 0.68 % |
| Number of trials analyzed | 9952 | 7212 | 5191 |

**Table 3.**
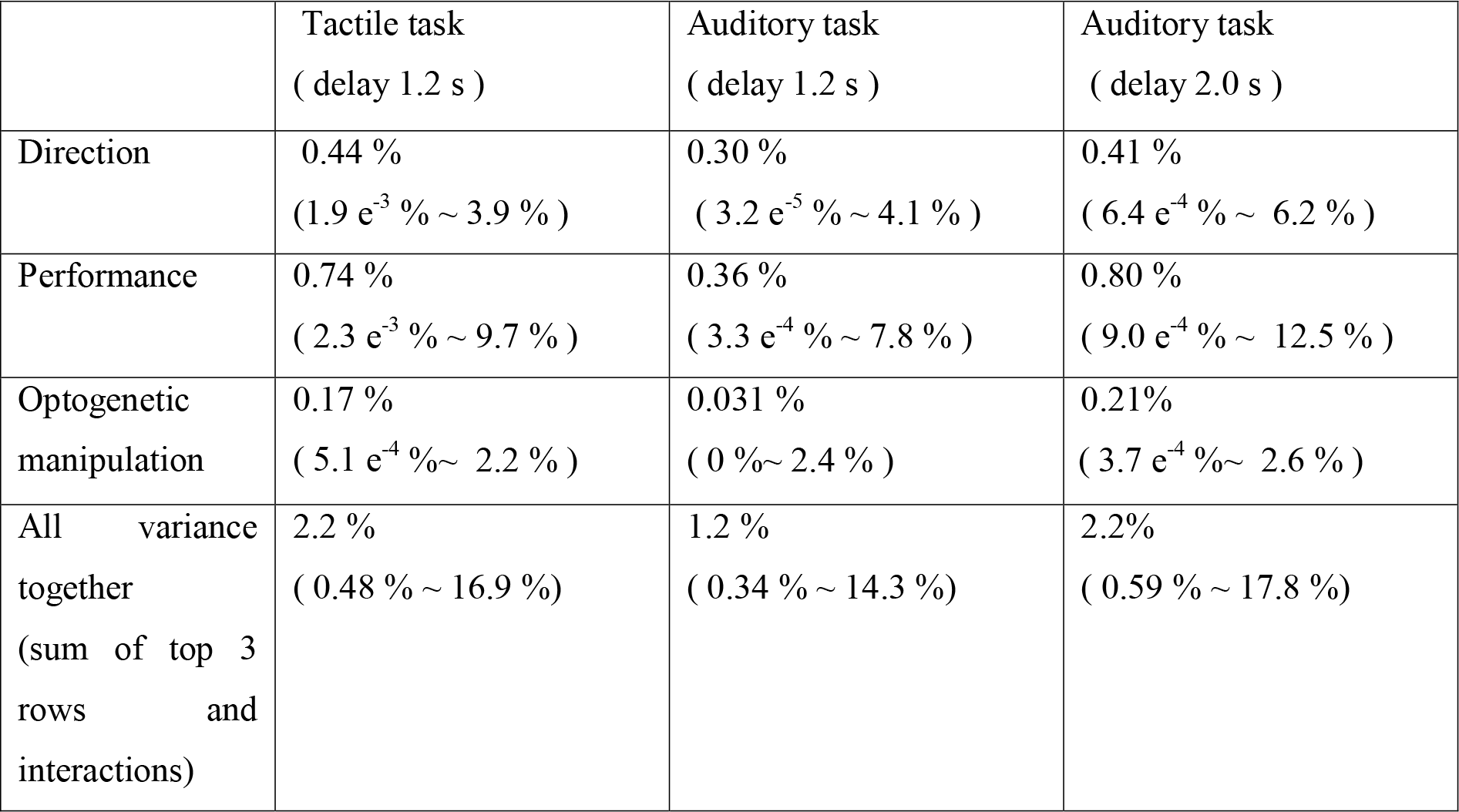
Variance of spike rate explained by previous trial types. Variance of spike rates in the pre-sample epoch explained by previous trial. We performed multifactor two-way ANOVA to test whether direction, performance, optogenetic manipulation, and their interactions affected the spike rate in each pyramidal cell. The median (top) and 95 % range (bottom, 2.5 % ~ 97.5 %) of the variance explained are shown in the table. The remaining variance is unexplained by previous trial type. Consistent with the effect of previous trials on behavior (Table 2), performance of previous trial affected the spike rates the most, while the optogenetic manipulation affected the spike rates the least.

### ALM is required for motor planning

The anterior lateral motor cortex (ALM; anterior 2.5mm, lateral 1.5mm to bregma) ^36^ is required for motor planning in the tactile delayed response task ^8^. To identify the dorsal neocortical regions required for motor planning in the auditory delayed response task, we performed an inactivation screen ^8^. Small parts of cortex (approximate radius, 1mm) ^8^ were ‘photoinhibited’ by photostimulating channelrhodopsin-2 (ChR2) expressed in parvalbumin (PV) positive interneurons (PV-ires-cre ^37^×Ai32 ^38^ animals). We photoinhibited 48 sites during the sample or delay epochs (Fig. 2a). Photoinhibiting the anterior frontal cortex unilaterally during the delay epoch biased future licking responses in the ipsilateral direction (Fig. 2b). The cortical area that caused a directional bias when silenced is indistinguishable from ALM ^8^. We did not detect any dorsal cortical region as required during the sample epoch. We confirmed the involvement of ALM in five additional mice (Fig. 2c, Table 4). Moreover, bilateral inactivation of ALM during the sample or delay epoch resulted in near chance level performance (Fig. 2d) ^30^. Bilateral inactivation in nearby cortical regions (posterior to ALM by 2mm) did not affect performance. The fact that even sample epoch inactivation resulted in chance level performance implies that other brain regions cannot rescue motor planning after ALM silencing. Finally, bilateral silencing during the first or second half of the 2 s delay epoch produced chance level performance {Inagaki et al, *in prep*}. Altogether, ALM is necessary for motor planning in a delayed-response licking task regardless of sensory modality and delay duration.

**Figure 2.**
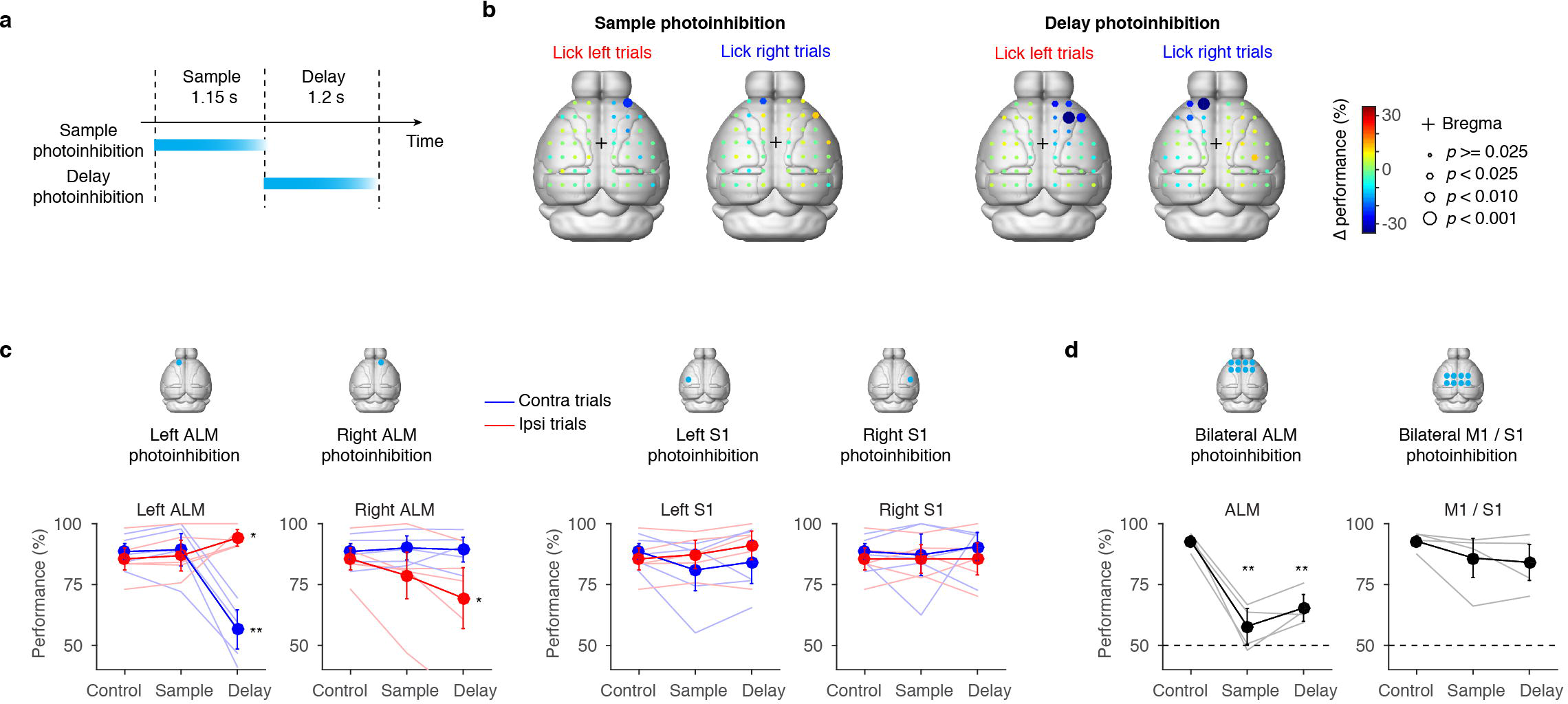
ALM is required for motor-planning. **a.** Schematic, photoinhibition of ALM either during the sample epoch or the delay epoch. Cortical regions were photoinhibited for 1.2 s starting at the onset of the sample or delay epochs (relevant to **b-d**). The photostimulus was ramped down over the last 200 ms to avoid rebound activity ^8^. **b.** Spatial maps of behavioral effects caused by photoinhibition during the sample epoch (left) and the delay epoch (right). Performance compared to the control condition Δperformance) is shown in color code. Spot sizes indicate *p*-values based on *hierarchical bootstrap* (Methods). *p*-values lower than 0.001 were significant even after correction for multiple comparisons (*Benjamini-Hochberg procedure*). Data based on 3 animals, 104 sessions; 50594 trials (75.1 ± 9.5 stimulation trials per spot, mean ± s.d.) for the sample photoinhibition; 50585 trials (101.2 ± 10.8 stimulation trials per spot, mean ± s.d.) for the delay photoinhibition. Control performance was 83.0 ± 4.0 (mean ± s.d.) % for contra trials, 85.1 ± 0.9 (mean ± s.d.) % for ipsi trials. **c.** Behavioral effects of unilateral ALM photoinhibition. Top, locations of photoinhibition. Bottom, behavioral performance. Thick lines, grand mean performance (n = 5 animals); thin lines, mean performance for each animal. Error bar, S.E.M. based on *hierarchical bootstrap* (Methods); Sample, trials with photoinhibition in the sample epoch; Delay, trials with photoinhibition in the delay epoch. The whisker representation area of the primary somatosensory cortex (S1) was photoinhibited as a control (see Methods for coordinates). Data based on 5 animals, 47 sessions, 12556 control trials and 3385 stimulation trials in total. *p*-values are based on *hierarchical bootstrap.* **, *p* < 0.001; *, *p* < 0.05 (both without correction for multiple comparisons. Only ** have *p* < 0.05 after *Bonferroni* correction). Fig. 2d follows the same format. See table 4 for *p*-values. **d.** Behavioral effects of bilateral ALM photoinhibition. Eight spots surrounding ALM in both hemispheres were photoinhibited (Methods). M1 was inhibited as a control (see Methods for coordinate). Data based on 4 animals, 40 sessions, 6096 control trials and 1125 stimulation trials in total. From left to right, *p*-values < 0.0001, 0.0001, and *p* = 0.0767, 0.0736 (*hierarchical bootstrap*, 10000 iterations, without correction for multiple comparison).

**Table 4.** *p*-values in Fig.2c. *p*-values are based on *hierarchical bootstrap*, 10000 iterations, without correction for multiple comparison (Methods).

| Stim location | Left ALM |  | Right ALM |  | Left S1 |  | Right S1 |  |
| --- | --- | --- | --- | --- | --- | --- | --- | --- |
| Epoch | Sample | Delay | Sample | Delay | Sample | Delay | Sample | Delay |
| Left lick trial | 0.3440 | 0.0196 | 0.1116 | 0.0147 | 0.3170 | 0.0786 | 0.5244 | 0.5369 |
| Right lick trial | 0.6489 | 0.0000 | 0.3057 | 0.4130 | 0.1064 | 0.3061 | 0.4802 | 0.3036 |

### Delay activity in ALM

We recorded single units from ALM using high-density silicon probes (See Table 1 for number of animals, sessions, and units) (Fig. 3, 4). All recordings were performed in the left ALM. Therefore, contralateral and ipsilateral directions refer to right and left licking, respectively. Correct lick right or left trials are referred to as “contra trials”(blue) or “ipsi trials” (red), respectively. We classified units into regular-spiking and fast-spiking neurons units based on their spike widths (Methods) ^8^. Here we focus on regular spiking units, or putative pyramidal neurons, unless otherwise described.

**Figure 3.**
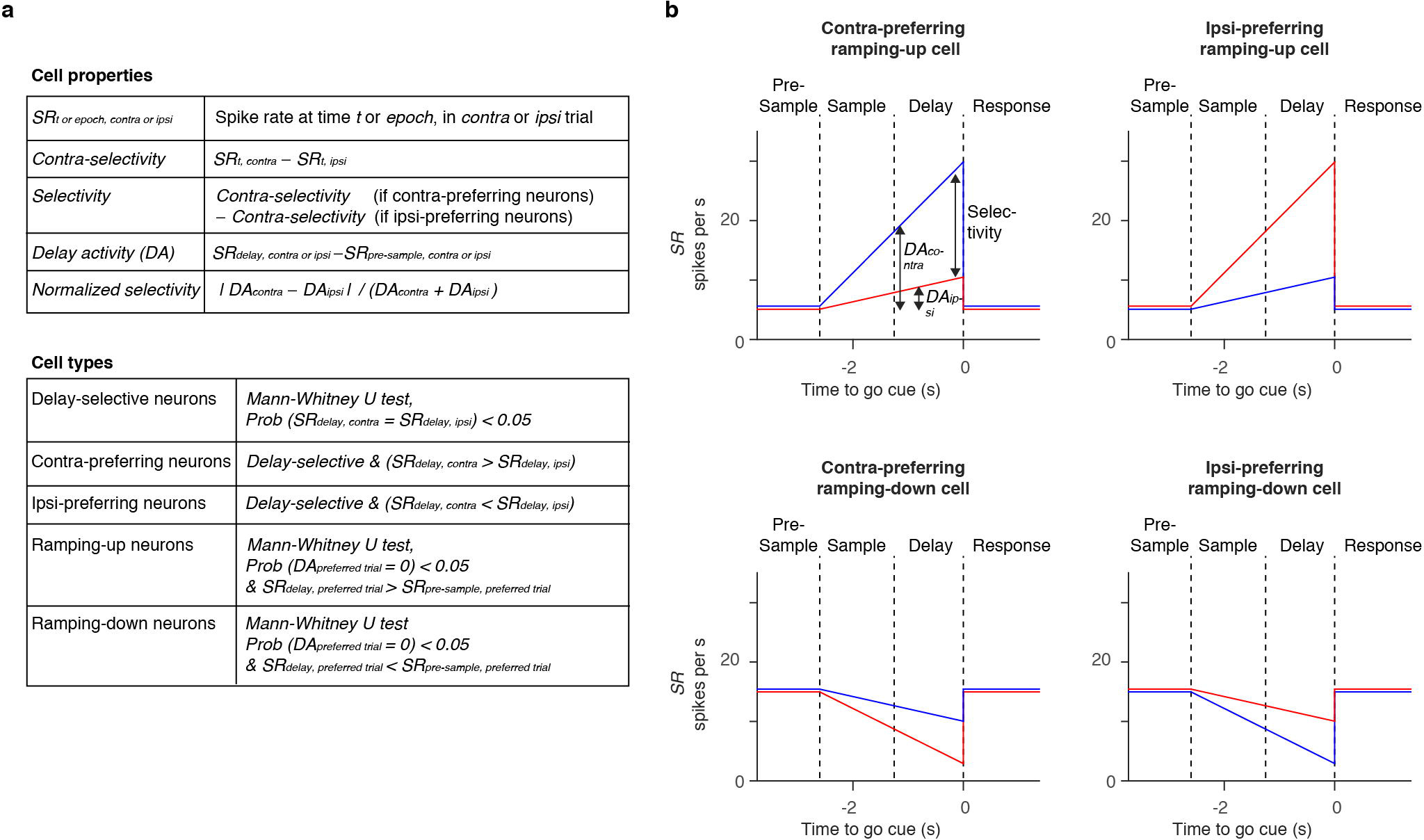
Classification of neural activity patterns. **a.** Parameters used to classify activity patterns. **b.** Schematics illustrating activity patterns. Blue, spike rates in contra trials. Red, spike rates in ipsi trials.

**Figure 4.**
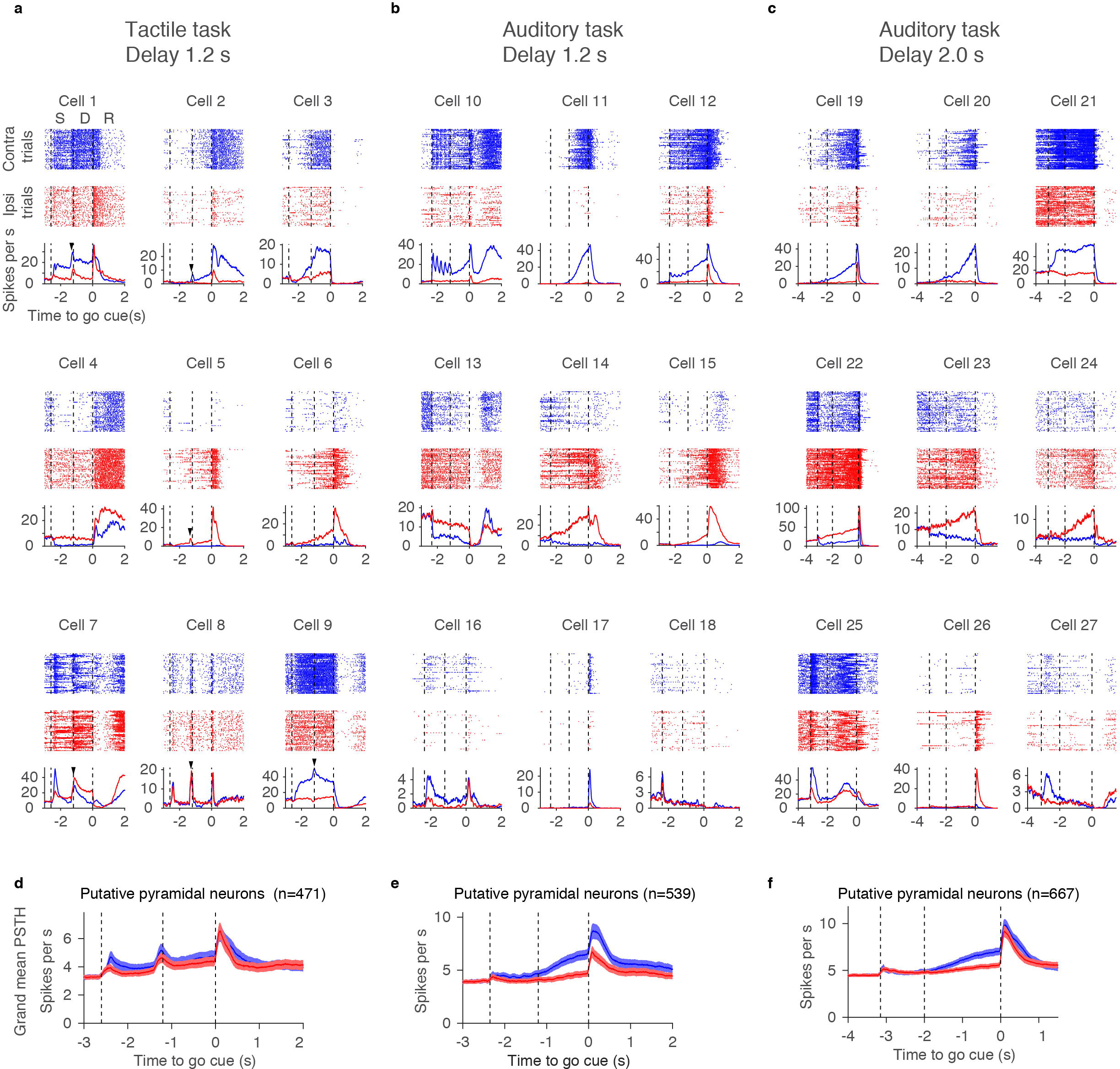
Example neurons in ALM. **a-c.** Nine example ALM neurons for each task type. Top, spike rasters. Bottom, peri-stimulus time histogram (PSTH). Blue, correct lick right trials (contra trials); red, correct lick left trials (ipsi trials). Randomly selected 50 trials are shown per trial type. Dashed lines separate behavioral epochs. S, sample epoch; D, delay epoch; R, response epoch. Time is relative to the go cue. Arrowheads indicate phasic activity at the beginning of the delay epoch. **d-f.** Grand mean PSTH for all pyramidal cells. Shadow, S.E.M. (*bootstrap*).

The majority of putative pyramidal neurons (88 %, Table 1) showed significant selectivity (trial type difference in spike count, *Mann*–*Whitney U* test, *p* < 0.05) in one or more of the three epochs (sample, delay, response epochs) (Fig. 5a-f). Proportions of selective neurons were lowest during the sample epoch, and highest during the response epoch (Fig. 5d-f). The majority of selective cells showed both ‘pre-movement’ and ‘peri-movement’ selectivity (Fig. 5a-c, bar). Some cells switched the sign of their selectivity between epochs (less than 20 % under all conditions) (Fig. 5g-i).

**Figure 5.**
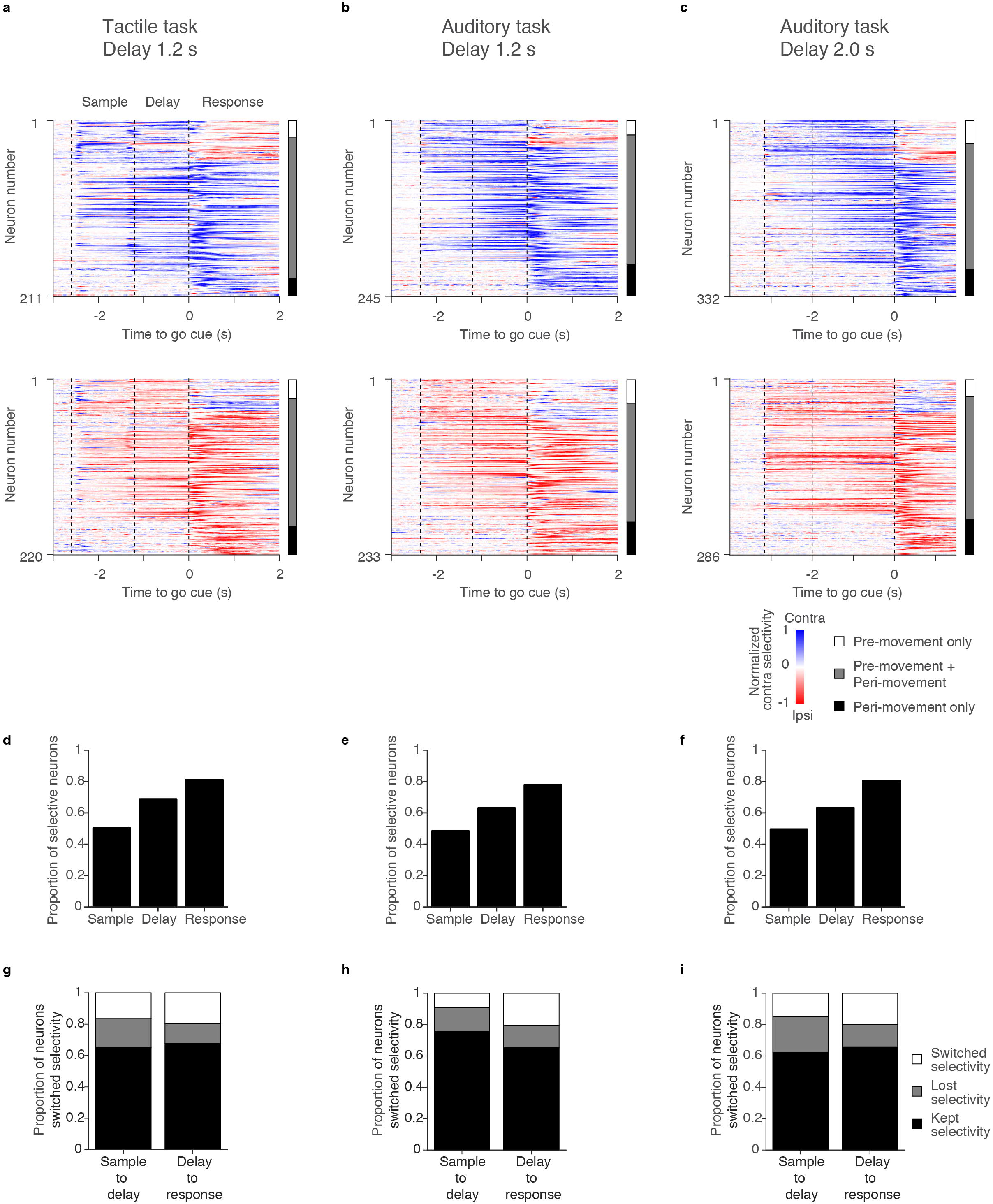
Selectivity in ALM. **a-b.** ALM population selectivity. Top, cells with higher spike counts across all trial epochs in the contra trials; Bottom, cells with higher spike counts in the ipsi trials. Each row represents contra selectivity normalized by the peak selectivity of individual neuron (blue, contra; red, ipsi). Vertical bars on the right; white, neurons with pre-movement (sample or delay epoch) selectivity only; grey, both pre-movement and peri-movement (response epoch) selectivity; black, peri-movement selectivity. **d-f.** Proportion of neurons with selectivity in each epoch. **g-i.** Proportion of neurons that kept, lost, or switched selectivity between epochs.

Sample epoch activity differed across the behavioral tasks (Fig. 4a-c). In the tactile task, many neurons showed a phasic signal at the transition from sample to delay epoch (Arrowheads in Fig. 4a). The grand mean spike rate of putative pyramidal neurons also showed this phasic response (Fig. 4d). This response is likely caused by a transient increase in whisker movement and touch input at the end of the sample epoch ^8,28^. In contrast, in the auditory task, some neurons showed spike rate changes that were phase-locked to the five repeated presentations of the tones (e.g. Fig. 4b, cell 10).

During the delay epoch, neural dynamics were diverse within a task (Fig. 4a-c), but the types of dynamics were similar across tasks (Fig. 4a-c). Many cells showed ramping-up activity during the delay epoch on contra trials (Fig. 4a-c, cell 2, 11, 12, 19, and 20) or ipsi trials (cell 5, 6, 14, 15, 23, and 24). Other cells showed ramping-down or suppression during the delay epoch, mainly in contra trials (cell 4, 13, 14, and 23). Some non-selective cells showed small changes in activity during the delay epoch (cell 18 and 27). In the rest of this paper we focus on delay epoch activity, because this activity is necessary for motor planning (Fig. 2).

More than 63 % of pyramidal cells showed significant selectivity during the delay epoch (“delay-selective cells”, Table 1, Fig. 3a, Fig. 6a-c). We refer to cells with significantly higher spike rate (*SR*) during the delay epoch of the contra trials as “contra-preferring neurons” and vice versa (*Mann–Whitney U* test, *p* < 0.05, Fig. 3a). In all tasks, “contra-preferring neurons” and “ipsi-preferring neurons” were spatially intermingled within a hemisphere (Fig. 6a-c, top vs. middle), consistent with previous reports in ALM and other brain regions and species ^4,9,29^. Although we classified cells into three classes, the distribution of selectivity was continuous (Fig. 6g-i). The selectivity of contra-or ipsi-preferring cells ramped to the beginning of the response epoch (Fig. 6d-f). On average, contra-preferring neurons ramped up during the contra trials, with little modulation during the ipsi trials (Fig. 6a-c, top). In contrast, ipis-preferring neurons ramped down during the contra trials with modest modulation during the ipsi trials (Fig. 6a-c, middle).

**Figure 6.**
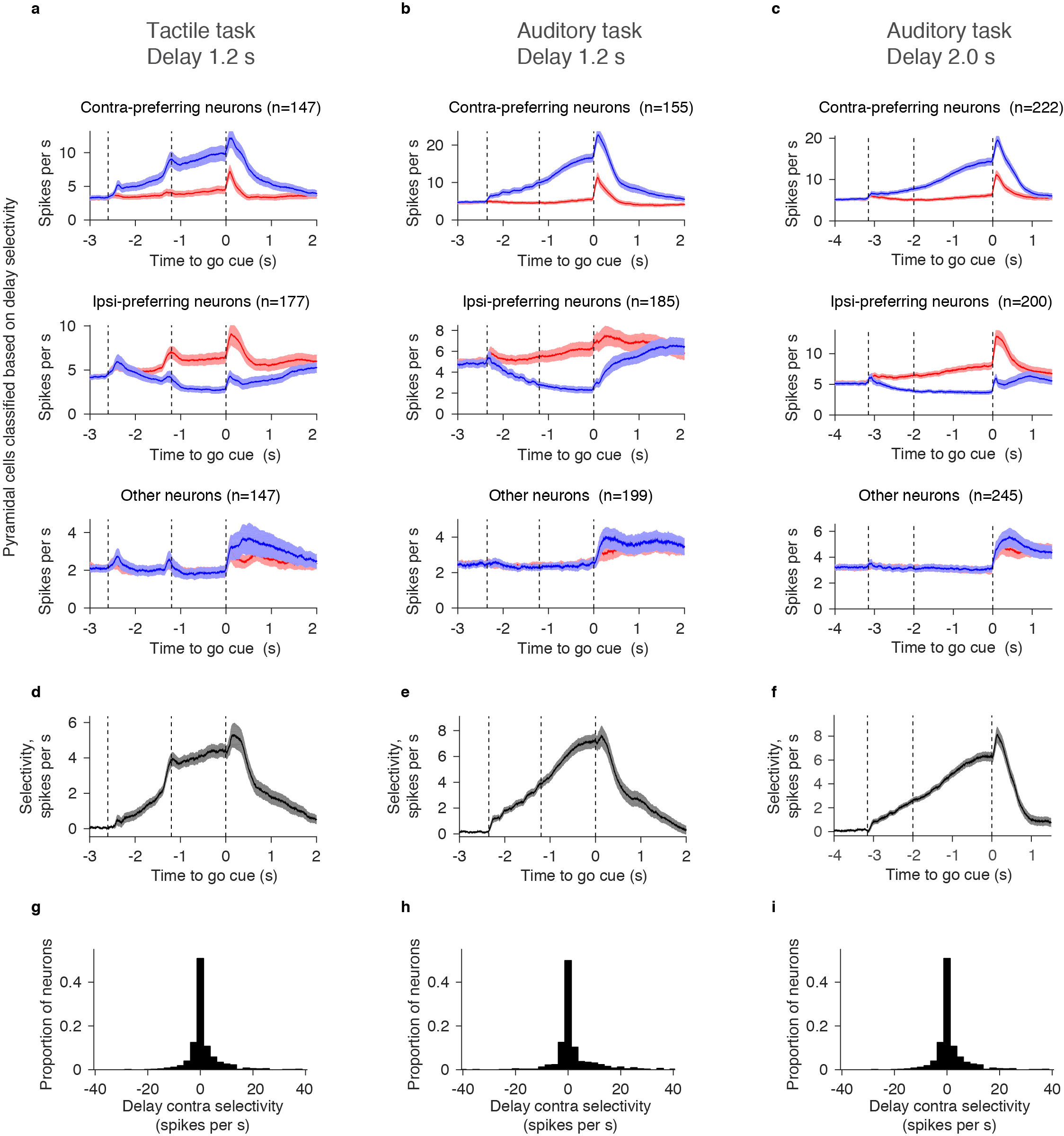
Selectivity and mean PSTH. **a-c.** Grand mean PSTH for cells with different types of delay selectivity. Shadow, S.E.M. (*bootstrap*). **d-f.** Grand mean selectivity for cells with significant delay selectivity (Methods). Shadow, S.E.M. (*bootstrap*). **g-i.** Distribution of delay contra selectivity. Bin size, 2.5 spikes per s.

‘Ramping-up’ (‘ramping-down’) cells have higher (lower) spike rates during the delay epoch compared to the pre-sample baseline in the preferred trial type (*Mann–Whitney U* test, *p* < 0.05, Fig. 3a, b). Contra-preferring neurons were mostly ramping-up cells during the delay epoch (Fig. 7a-f). In contrast, ipsi-preferring neurons were either ramping-up or down (Fig. 7a-c). The latter were classified as ipsi-preferring because the ramping-down was stronger in contra trials. Because of this asymmetry, overall, there were more contra ramping-up cells than ipsi ramping-up cells for all tasks (Fig. 7d-f).

**Figure 7.**
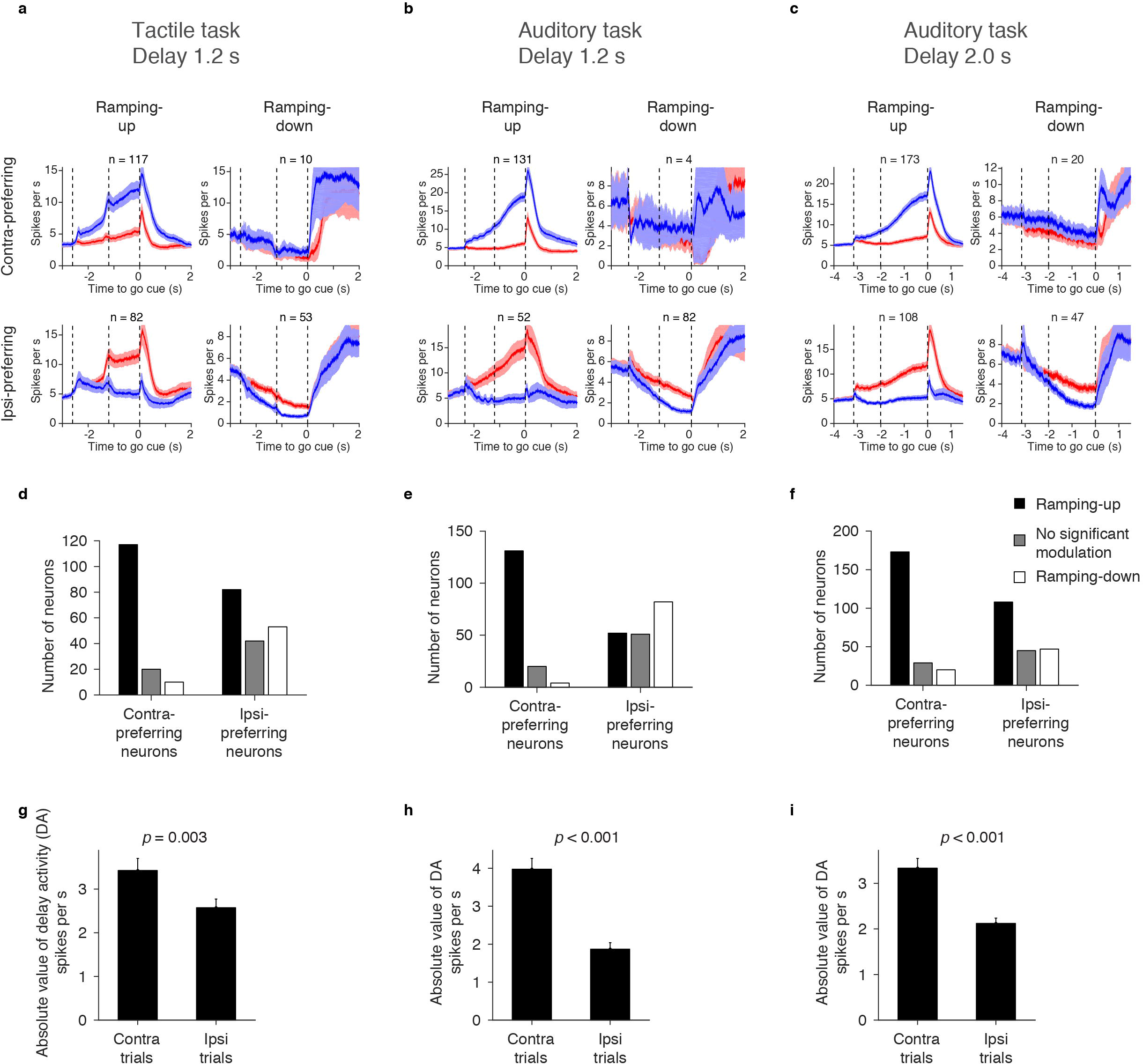
Selectivity and ramping. **a-c.** Grand mean PSTH of contra-preferring and ipsi-preferring cells with different ramping directions. Ramping up/down was tested in the preferred direction (*Mann–Whitney U test*, *p*< 0.05). Shadow, S.E.M. (*bootstrap*). **d-f.** Numbers of neurons with different ramping directions. **g-i.** Comparison of the absolute value of delay activity (DA) (Methods) between the contra and ipsi trials. Mean ± S.E.M. (*bootstrap*) is shown in the bar graph. All putative pyramidal cells were included. *P*-value is based on *bootstrap* with 1000 iterations testing a null hypothesis that contra DA is lower than ipsi DA.

We next characterized lateralization of selectivity. We defined “delay activity (DA)” in each cell as the spike rate difference between the pre-sample baseline epoch and the delay epoch (*SR*_*delay*_–*SR*_*pre-sample*_)_*trial type*_, where ‘trial type’ is either contra or ipsi, and *SR* denotes mean spike rate during the delay or pre-sample epoch in each trial type, Fig. 3a). The absolute value of the DA was higher on contra trials compared to ipsi trials (Fig. 7g-i), as were the number of contra ramping-up neurons. This accounts for the contralateral bias in the grand mean peri-stimulus time histogram (PSTH) (Fig. 4e-f). In contrast, in the tactile task ramping-up and down cells canceled each other and the grand mean PSTH was similar for the two trial types (Fig. 4d). The contralateral selectivity in the auditory task is similar to activity in prefrontal cortex of primates during memory guided saccade tasks ^4^.

What is the relationship of DA across trial types? The contra-selective ramping-up cells did not decrease spike rates in ipsi trials, on average (Fig. 7a-c). In contrast, DA also increased slightly in the ipsi trials (Fig. 8a-c, blue lines, regression). Similarly, the ipsi-selective ramping-up cells showed only a slight activity change during the contra trials (Fig. 7a-c and Fig. 8a-c, red lines). Therefore, DA in ramping-up cells were nearly independent between the two trial types: cells that ramped-up in the preferred trials had only weak increase in the non-preferred trials.

**Figure 8.**
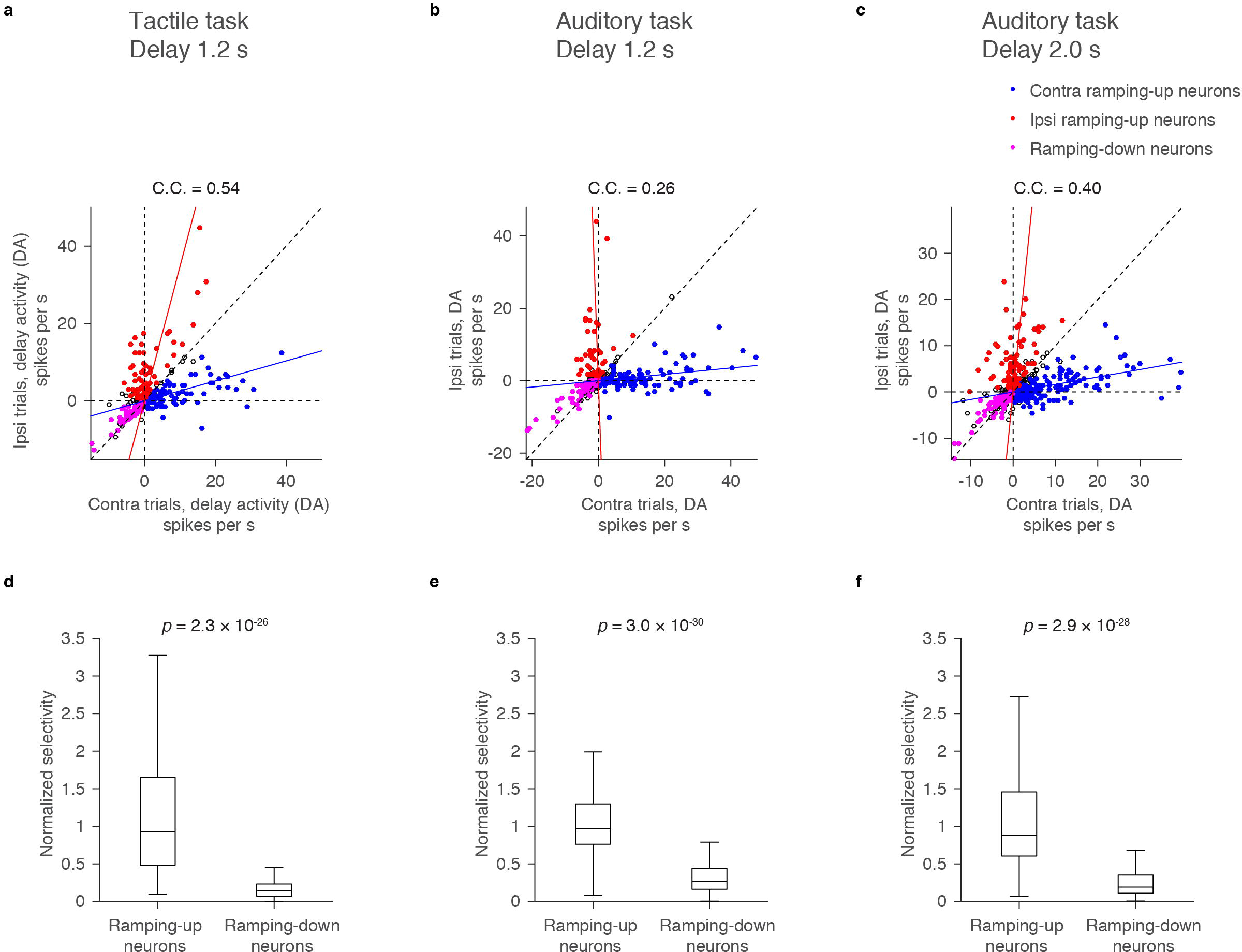
Relationship of delay activity (DA) in contra and ipsi trials. **a-c.** Distribution of contra and ipsi DA of all putative pyramidal neurons. Each circle corresponds to one neuron. Contra ramping-up cells, ipsi ramping-up cells, and ramping-down cells (both contra and ipsi ramping-down cells) are shown as filled circles in different color. Other cells are shown as open circles. Line represents linear regressions (blue, contra ramping-up cells; red, ispi-ramping-up cells). C.C, correlation coefficient of contra and ipsi DA based on all cells. Diagonal dotted line, line with slope one. **d-f.** Normalized selectivity (Methods) of ramping-up neurons (both contra-and ipis-ramping-up neurons) and ramping-down neurons (both contra-and ipis-ramping-down neurons). Central line in the box plot is the median. Top and bottom edges are the 75 % and 25 % points, respectively. The whiskers show the lowest datum within 1.5 interquartile range (IQR) of the lower quartile, and the highest datum within 1.5 IQR of the upper quartile. *P*-values, *Mann–Whitney U* test.

In contrast to the near orthogonal DA of ramping-up cells, both contra and ipsi ramping-down cells ramped down for both movement directions (Fig. 7a-c, and Fig. 8a-c, magenta). We defined the normalized selectivity as | *DA*_*contra*_ - *DA*_*ipsi*_ | / (*DA*_*contra*_ + *DA*_*ipsi*_) (Fig. 3a). Weaker normalized selectivity of the ramping-down cells compared to the ramping-up cells (Fig. 8d-f) is consistent with recordings in prefrontal cortex in non-human primates ^4^.

Similarly, putative fast-spiking GABAergic interneurons (FS neurons) ^8^ had weaker normalized selectivity than putative pyramidal cells (Fig. 9g-l), while they had strong absolute selectivity because of high spike rates (Fig. 9a-f). Consistently, correlations between contra and ipsi DA were higher in FS neurons (Fig. 8a-c vs. Fig. 9g-i). Altogether, both ramping-down cells and FS neurons showed similar activities between two trial types. These findings are consistent with FS neurons pooling the activity of local pyramidal neurons ^39^ and inhibiting the ramping-down cells.

**Figure 9.**
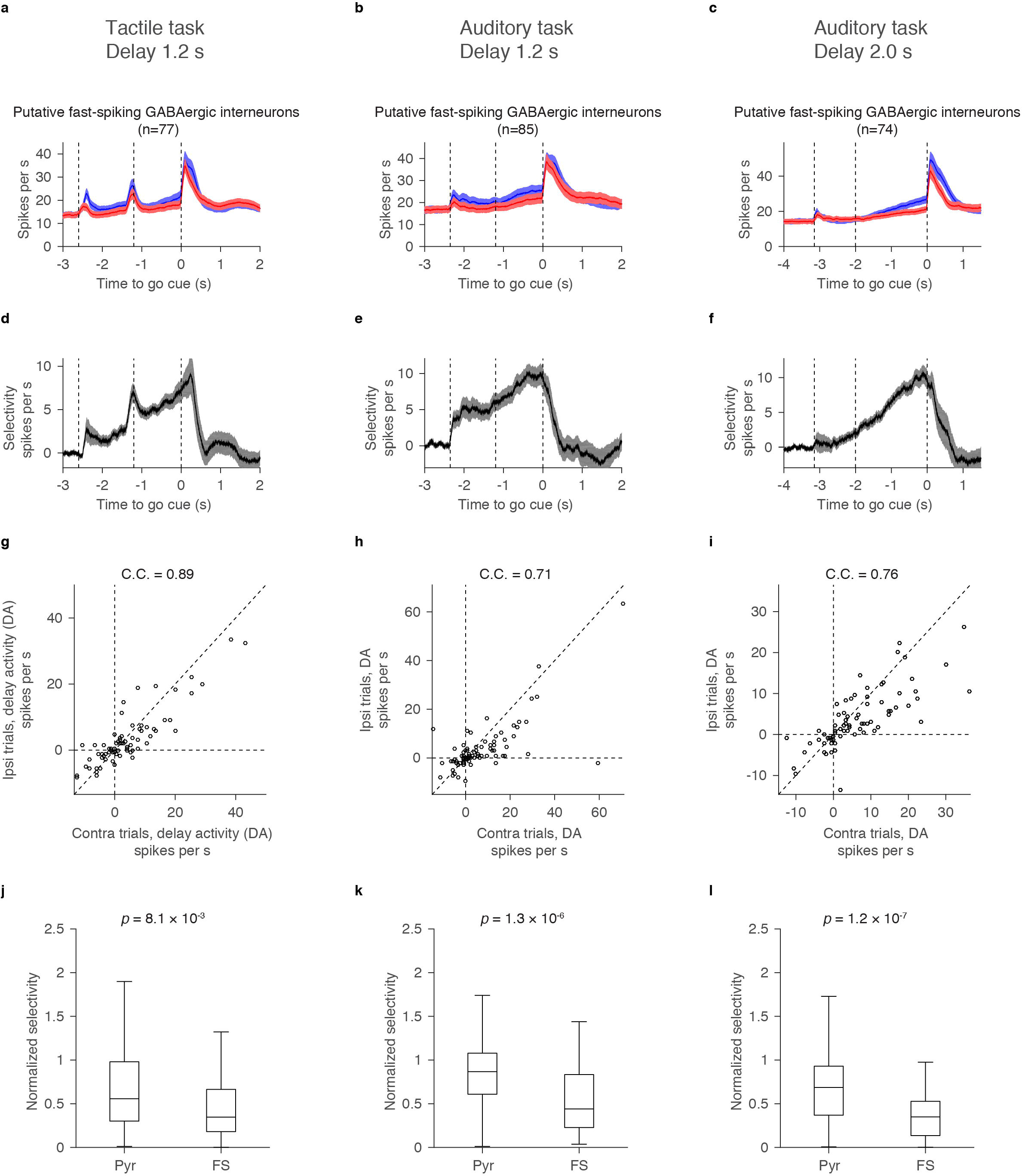
Selectivity of putative fast-spiking GABAergic interneurons (FS neurons) **a-c.** Grand mean PSTH for all FS neurons. Shadow, S.E.M. (*bootstrap*). **d-f.** Grand mean selectivity of FS neurons with significant delay selectivity. Shadow, S.E.M. (*bootstrap*). **g-i.** Distribution of contra and ipsi DA for all FS neurons. Each circle corresponds to each cell. C.C, correlation coefficient of contra and ipsi DA. Diagonal dashed line, line with slope one. **j-l.** Normalized selectivity of putative pyramidal neurons (pyr) and FS neurons. For these figures, normalized selectivity was measured for neurons with positive DA ( *DA*_*contra*_ + *DA*_*ipsi*_ > 0). Central line in the box plot is the median. Top and bottom edges are the 75 % and 25 % points, respectively. The whiskers show the lowest datum within 1.5 interquartile range (IQR) of the lower quartile, and the highest datum within 1.5 IQR of the upper quartile. *P*-values, *Mann–Whitney U* test.

### Preparatory activity in ALM

Does delay activity maintain information about the sensory input, or is it related to future movements, as required for preparatory activity? Calcium imaging and extracellular recordings have reported ALM neurons with preparatory activity in the tactile task ^8,28^. To confirm this finding in the auditory task we analyzed incorrect trials. Since incorrect trials were rare in the cohort of mice trained on the auditory task with 1.2 s delay (performance, 91.4 %), we compared the tactile task and the auditory task with 2.0 s delay (performance, 80.3 % and 87.2 %, respectively). We analyzed neurons with delay selectivity that were recorded in sessions with at least ten incorrect trials for both trial types (incorrect lick right and incorrect lick left trials) (tactile task, 412 cells; auditory task, 465 cells).

For each cell we compared selectivity during the second half of the delay epoch in correct and incorrect trials (Fig. 10a, b). Cells on the line with slope one 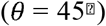 have the same selectivity in both correct and incorrect trials, indicating coding of the sensory stimulus (instruction). Cells on the line with slope negative one 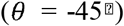 switch selectivity for correct and incorrect trials, indicating preparatory activity. The majority of cells had negative slopes 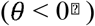, consistent with preparatory activity (Fig. 10c, d). Yet, *θ* was widely distributed, and the distributions were not centered on 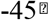 (Fig. 10c, d). Cells with slopes between one and negative one 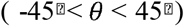 conjunctively code for sensory and motor information. Alternatively, these neurons may have lower selectivity in incorrect trials because of weaker perceived sensory input, attention, arousal, or confidence in incorrect trials ^40^. We did not detect a relationship between the distribution of and recording depth (Fig. 10e, f).

**Figure 10.**
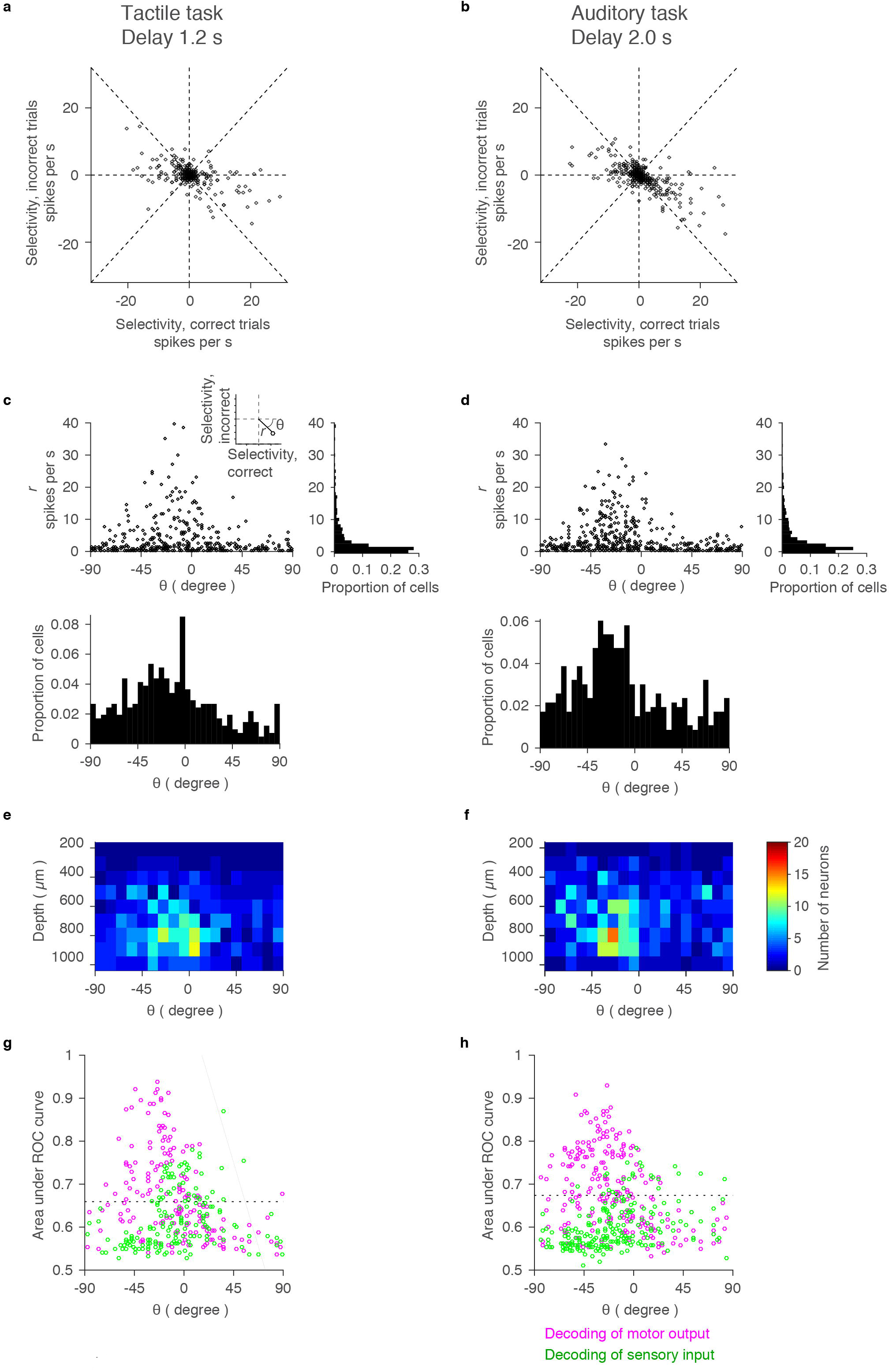
Coding during the delay epoch. **a-b.** Relationship between selectivity in correct and incorrect trials. We analyzed cells with more than 10 incorrect trials for each trial-type (incorrect lick-right and lick-left trials). Each point corresponds to each cell. Dashed lines: horizontal, vertical, and lines with slope one and minus one. **c-d.** Scatter plot and histogram of polar coordinates (*r* and *θ*) in **a-b**. Inset, the definition of *r* and *θ*. Bin size: 5 degrees for *θ*, 1 spikes per s for *r*. **e-f.** Distribution of *θ* across cortical depth. Bin size: 10 degrees for *θ*, 100 μm for depth. **g-h.** Decoding of motor and sensory information using ROC (Methods). *θ* and area under ROC curve for motor output (magenta) and sensory input (green) are shown for each cell. Dashed line, 95 % confident interval of shuffled data (Methods). Cells that have proportion correct higher than the dashed line were judged as cells decoding sensory or motor information.

Consistent with the polar coordinate analysis (Fig. 10a-d), many ALM neurons decoded motor output on a trial-by-trial basis (Area under receiver-operating characteristic (ROC) curve higher than the chance level. 85, 117 cells for the tactile and auditory task, respectively) (Fig. 10g, h, magenta above dashed line). In contrast, decoding of sensory information was relatively poor (45, 24 cells for the tactile and auditory task, respectively) (Fig. 10g, h, green above dashed line).

How does coding change over time within a trial? We measured *θ* throughout the trial. In both tasks, some cells coded for sensory information during the sample epoch (Fig. 11a-d, 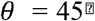). During the delay epoch, the number of neurons with negative *θ* gradually increased and peaked after the go cue. The distribution of *θ* was broad and did not have a peak at 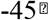. This analysis confirmed that throughout the delay epoch many cells show mixed coding with a bias toward preparatory activity.

**Figure 11.**
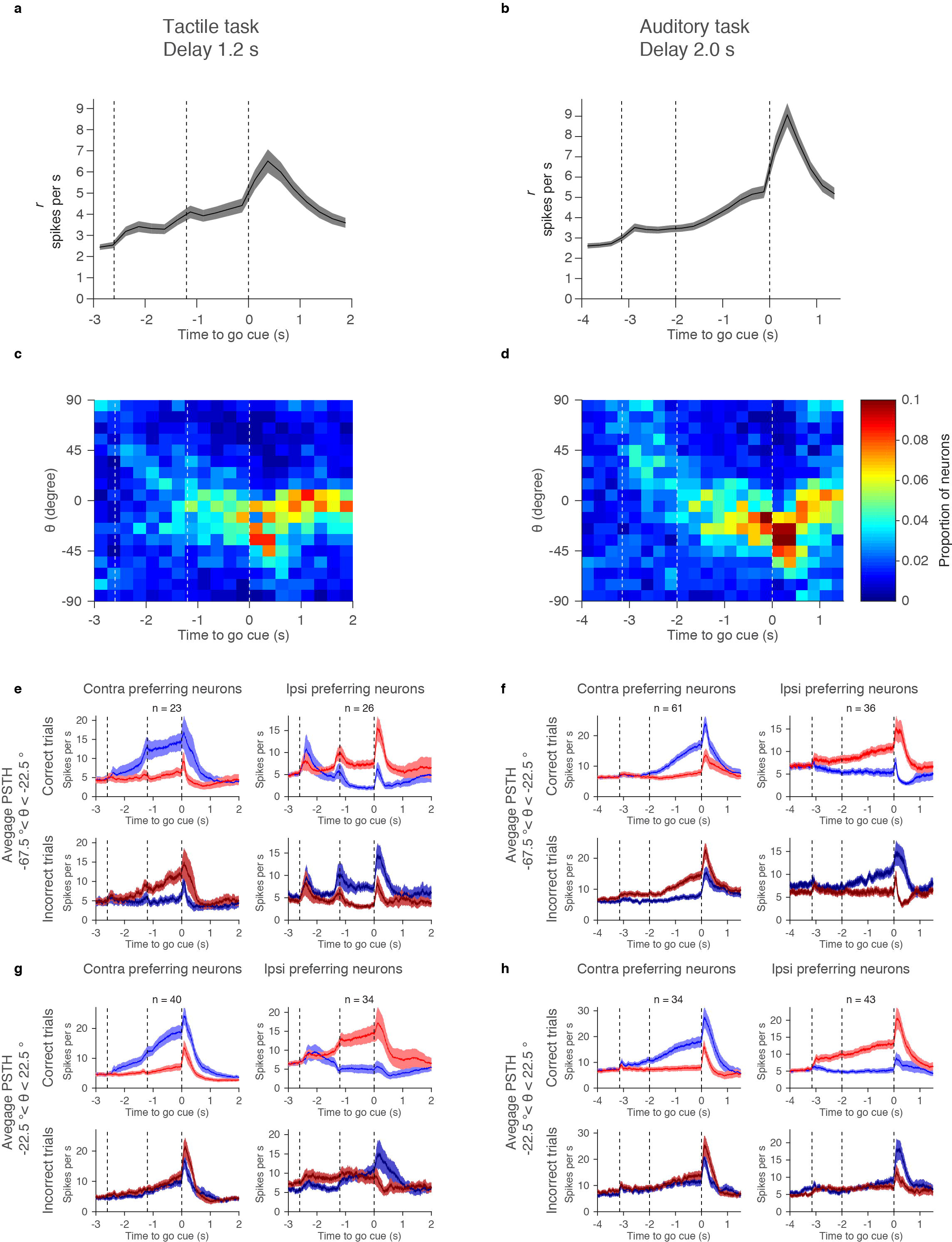
Coding of selectivity over time. **a-b.** Amplitude of selectivity (*r*) over time. Mean and S.E.M. (1000 permutations). **c-d.** Histogram of *θ* over time. Bin size: 10 degrees for *θ* and 250 ms for time. Mean of 1000 permutation. To calculate *θ*, we selected cells with *r* > 2 (Fig. 10a, b). Because of the ramping, the number of analyzed neurons increased during the delay epoch. **e-f.** Grand mean PSTH of cells coding motor-output 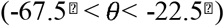. Shadow, S.E.M. (*bootstrap*). **g-h.** Grand mean PSTH of cells with mixed-selectivity coding 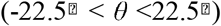. Shadow, S.E.M. (*bootstrap*).

Cells with preparatory activity 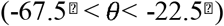 ramped from the sample epoch to the end of the delay epoch (Fig. 11e, f). Their selectivity flips between correct and incorrect trials, consistent with coding for motor-planning (Fig. 11e, f). Activity of mixed-selectivity neurons (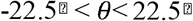) can be similar to preparatory cells in correct trials, but different in the incorrect trials (Fig. 11g, h top vs. Fig. 11e, f, top). This analysis confirms that preparatory activity can only be detected with reference to incorrect trials.

### Delay activity is monotonic and low-dimensional

So far we have shown that activity, averaged across cells, ramps during the delay epoch. This does not imply that each neuron also ramps (Fig. 12c). For instance, each neuron could have nonmonotonic activity peaking at different times in the trial, producing ramping on average (Fig. 12d) ^41^. Such sequences have been reported during motor planning and other kinds of short-term memory in premotor cortex, mPFC, PPC and hippocampus of rodents ^10,42-44^.

**Figure 12.**
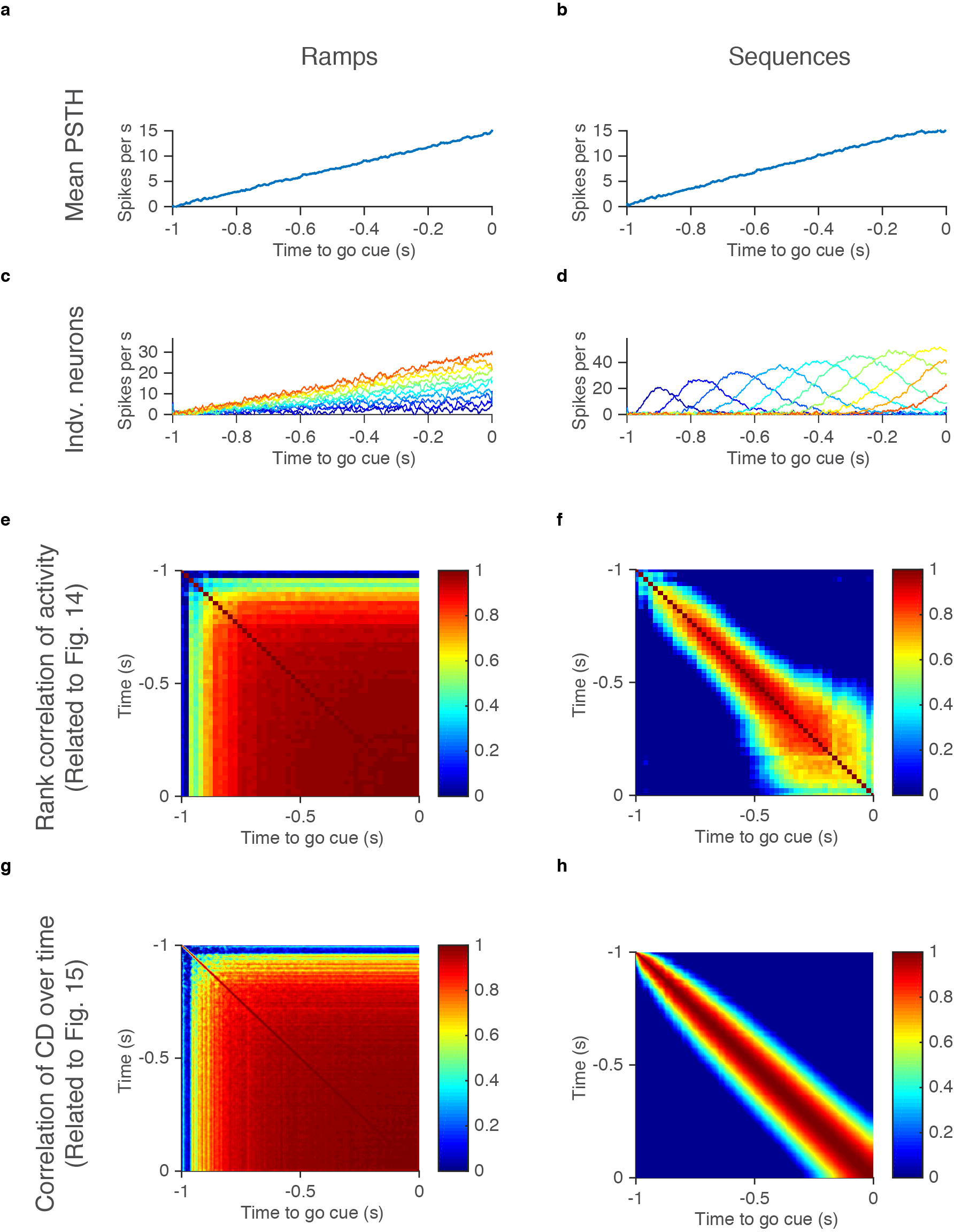
Two possible models producing ramping activity on average. **a-b.** Mean PSTH of all modeled cells in **c, d**. **c-d.** PSTH of individual modeled cells with ramping (**c**) or with bumps at different timing (**d**) (Methods). Eleven out of fifty modeled cells are shown. **e-f.** Rank correlation of neural activities over time (Related to Fig. 14). **g-h.** Stability of CD over time (Related to Fig. 15).

We performed four analyses to distinguish between these two scenarios (Fig. 12a vs. b). First, we tested if activity peaks are at the beginning or at the end of the delay epoch, as expected for ramping-down and up neurons. Alternatively, activity peaks could tile the entire delay epoch. For each delay-selective neuron, we selected a random set of trials and identified the peak in the average spike rate. The peaks tiled the delay epoch (Fig. 13a-c). We next calculated the mean spike rate based on the second half of trials and ordered them based on the rank order derived from the first half. The sequence of peaks disappeared, indicating that activity peaks were spurious (Fig. 13d-f). We repeated this analysis with 1000 permutations (Fig. 13g-l). The density of peaks was higher than chance level only at the beginning and at the end of the delay epoch, at the transition points between epochs (Fig. 13j-l). The cells with peaks at the beginning correspond to ramping-down cells and the cells with peaks at the end correspond to ramping-up cells. This analysis indicates that ALM cells are ramping.

**Figure 13.**
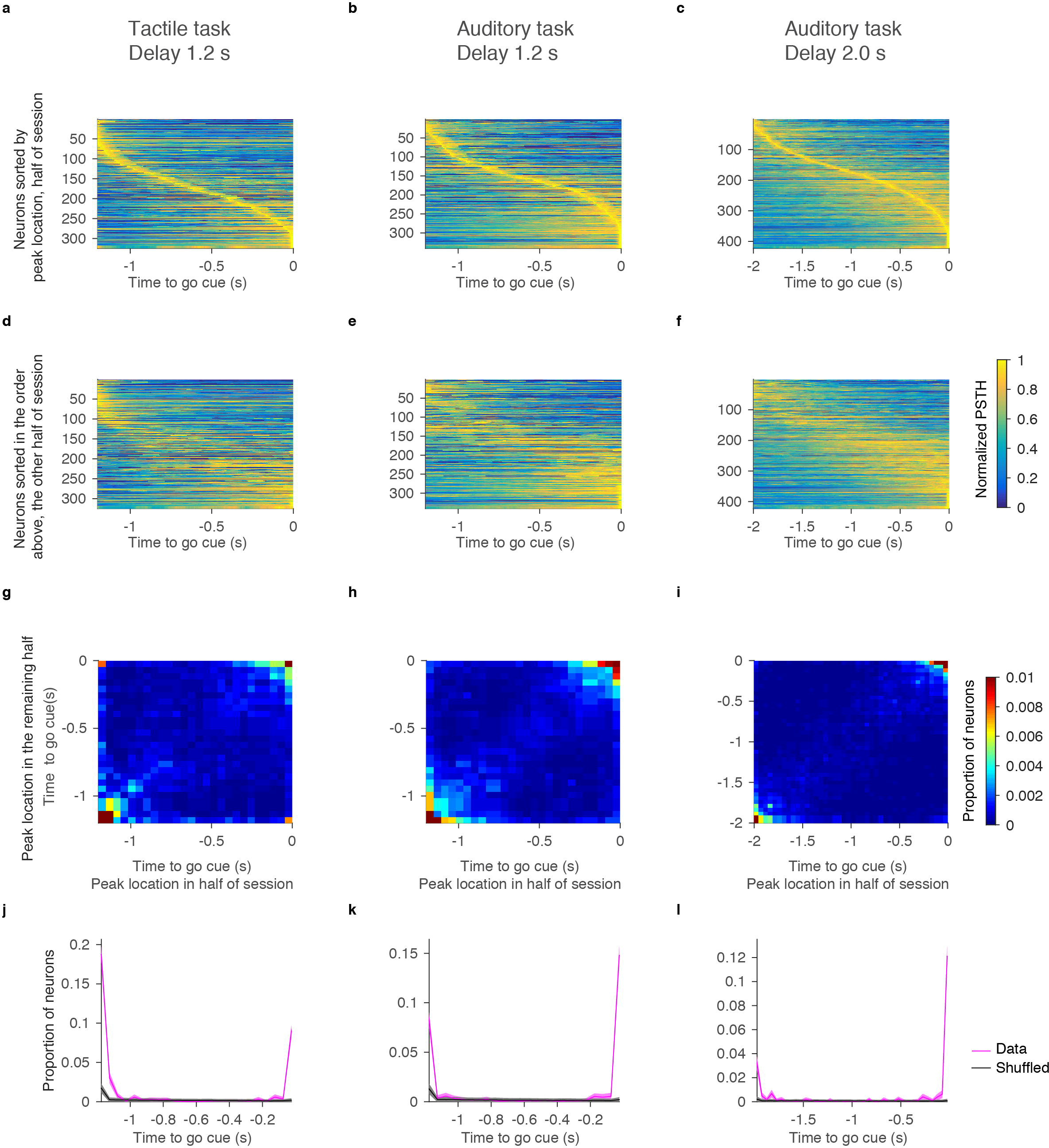
Activity peaks are at the beginning or end of the delay epoch. **a-c.** Location of activity peaks in randomly selected half of trials. Neurons were sorted by the activity peak location. Each row, PSTH of each cell normalized by its peak (colormap). **d-f.** Location of activity peaks in the remaining half of trials. Neurons were sorted in the same order as in **a-c**. **g-i.** Heatmaps indicating the density of peaks in the randomly selected half of trials and in the rest of trials (Methods). Bin size, 50ms. **j-l.** Proportion of neurons with peak at each time point (magenta). This corresponds to proportion of neurons on the positive diagonal axis in **g-i**. Black, chance level (Methods). Shadow, S.E.M. (bootstrap).

Second, if individual neurons ramp, their rank order should be approximately constant across time (Fig. 12c), and the Spearman’s rank correlation should be high (Fig. 12e). In contrast, if individual neurons are part of sequences, the rank correlation should be low across time (Fig. 12f). To exclude contributions of phasic activity at the transitions between sample and delay epochs, we analyzed the last 1.0 s of the 1.2 s delay (Fig. 14a, b), and 1.8 s of the 2.0 s delay (Fig. 14c). We plotted the spike rate change (SR_t_-SR_-1 s_ for 1.2 s delay, and SR_t_-SR_-1.8 s_ for 2 s delay) for all delay-selective neurons throughout this period (Fig. 14a-c). Each line represents spike rate change for one neuron. As noted above (Fig. 7g-i), ramping amplitudes were larger for contra trials than ipsi trials. Moreover, the Spearman’s rank correlation of spike rate change was high throughout the delay epoch (Fig. 14d-f). This implies that most cells with delay selectivity follow a similar time-course during the delay epoch and differ in terms of sign and amplitude. This is consistent with ALM ramping down or up during the delay epoch.

**Figure 14.**
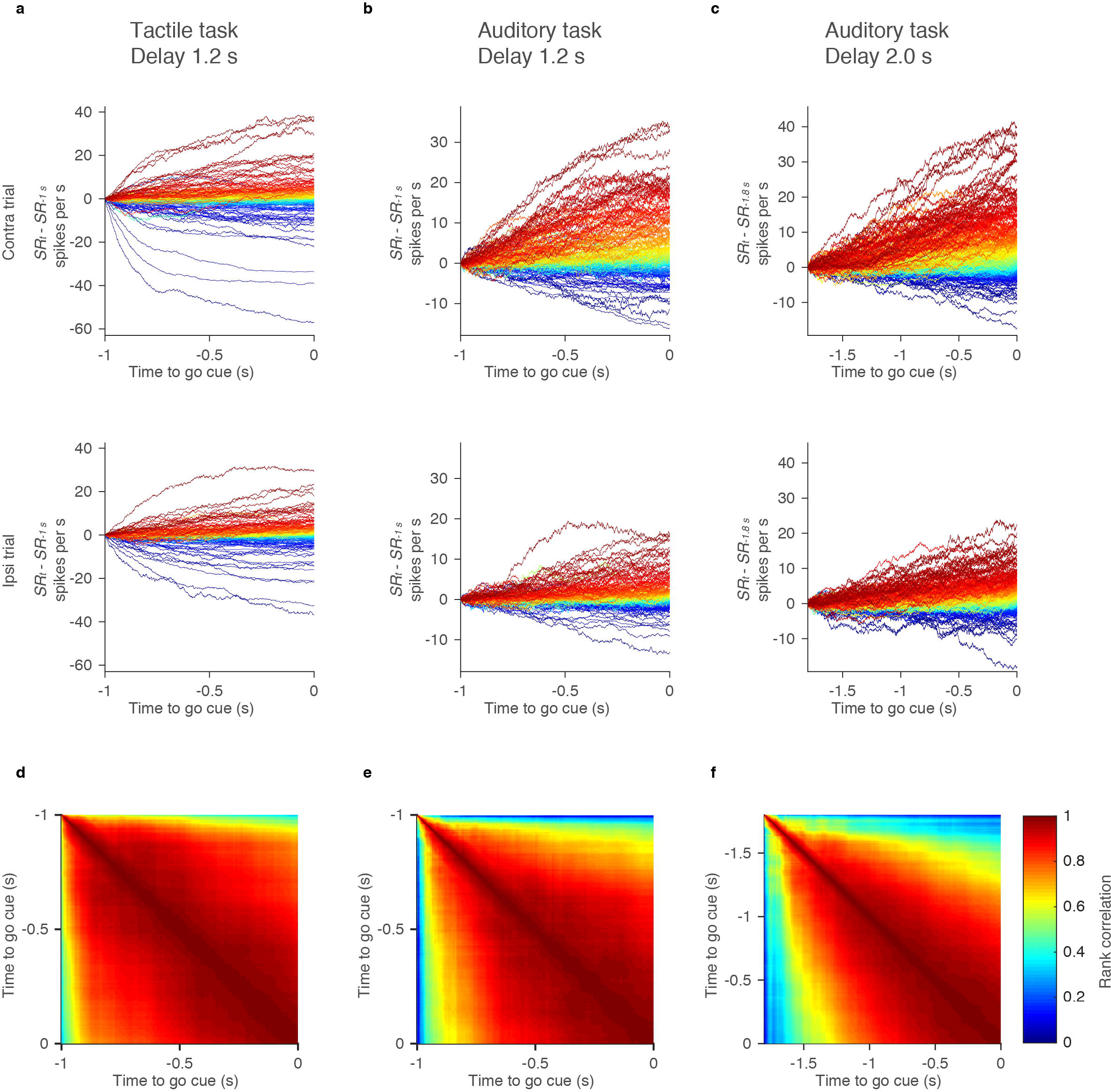
Rank correlation of delay activity (DA) **a-c.** Change in spike rate (averaged over 100ms) during the delay epoch in the contra trials (top) and ipsi trials (bottom). All cells with delay selectivity are shown. Each line, individual neuron. Lines were color coded according to rank order of change in spike rate at the end of the delay epoch. **d-f.** Rank correlation of change in spike rate between different time points.

Third, we analyzed the stability of decoders of trial types. The decoder is the direction in activity space that best distinguishes trial types (coding direction, CD) ^30^. CD is a vector defined in *n*-dimensional space, where *n* is the number of simultaneously recorded units (Fig. 15a). The CD should be stable for ramping activity because overlapping sets of neurons decode trial types throughout the delay epoch (Fig. 12g). In contrast, for sequences, CD should be unstable because different sets of neurons decode trial types at different time points (Fig. 12h).

**Figure 15.**
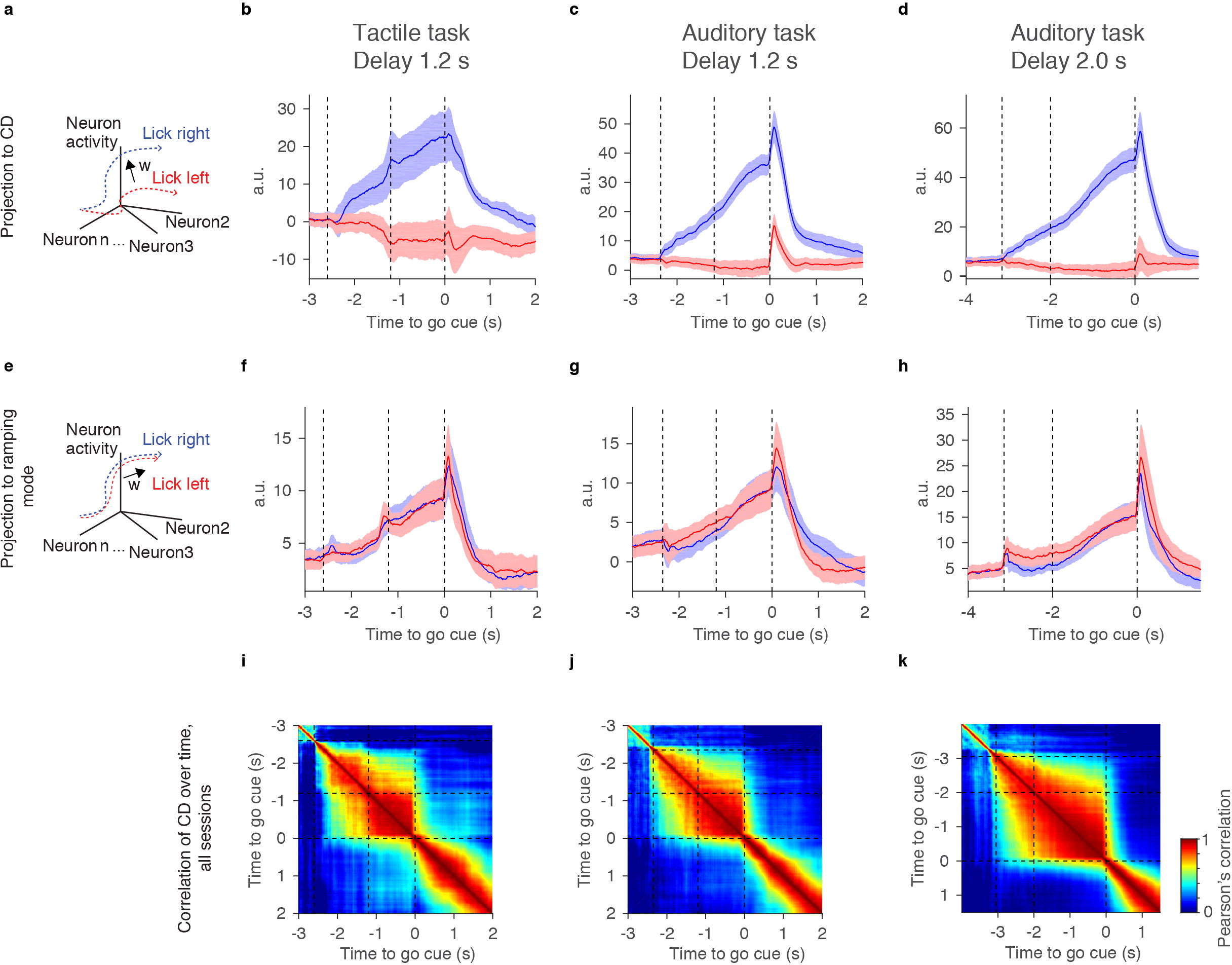
Stability of the coding direction (CD) **a.** Definition of CD as the direction that best differentiates trial type-related activity. **b-d.** Projection to the CD. Shadow, S.E.M. (*bootstrap*). **e.** The ramping mode (RM) is the first SVD mode capturing the largest remaining variance not explained by CD (Methods). It captures non-selective ramping as shown in **f-h**. **f-h.** Projection to the RM. Shadow, S.E.M. (*bootstrap*). **i-j.** Stability of CD. CD was calculated at each time point and compared with CD calculated at different time point using Pearson’s correlation (Methods).

We analyzed simultaneously recorded ALM neurons (see Fig. 16j-l and Table 1 for numbers of sessions and units per sessions). We estimated CD based on the spike rate difference between two trial types at each time point (Fig. 15a, Methods). The CD explained more than 90 % of variance of the selectivity (Fig. 16a-c, Methods). Moreover, the CD was stable throughout the delay epoch (Fig. 15i-k). This implies that the same set of neurons code trial type differences throughout the delay epoch, consistent with ramping activity, but inconsistent with sequences (Fig. 12).

**Figure 16.**
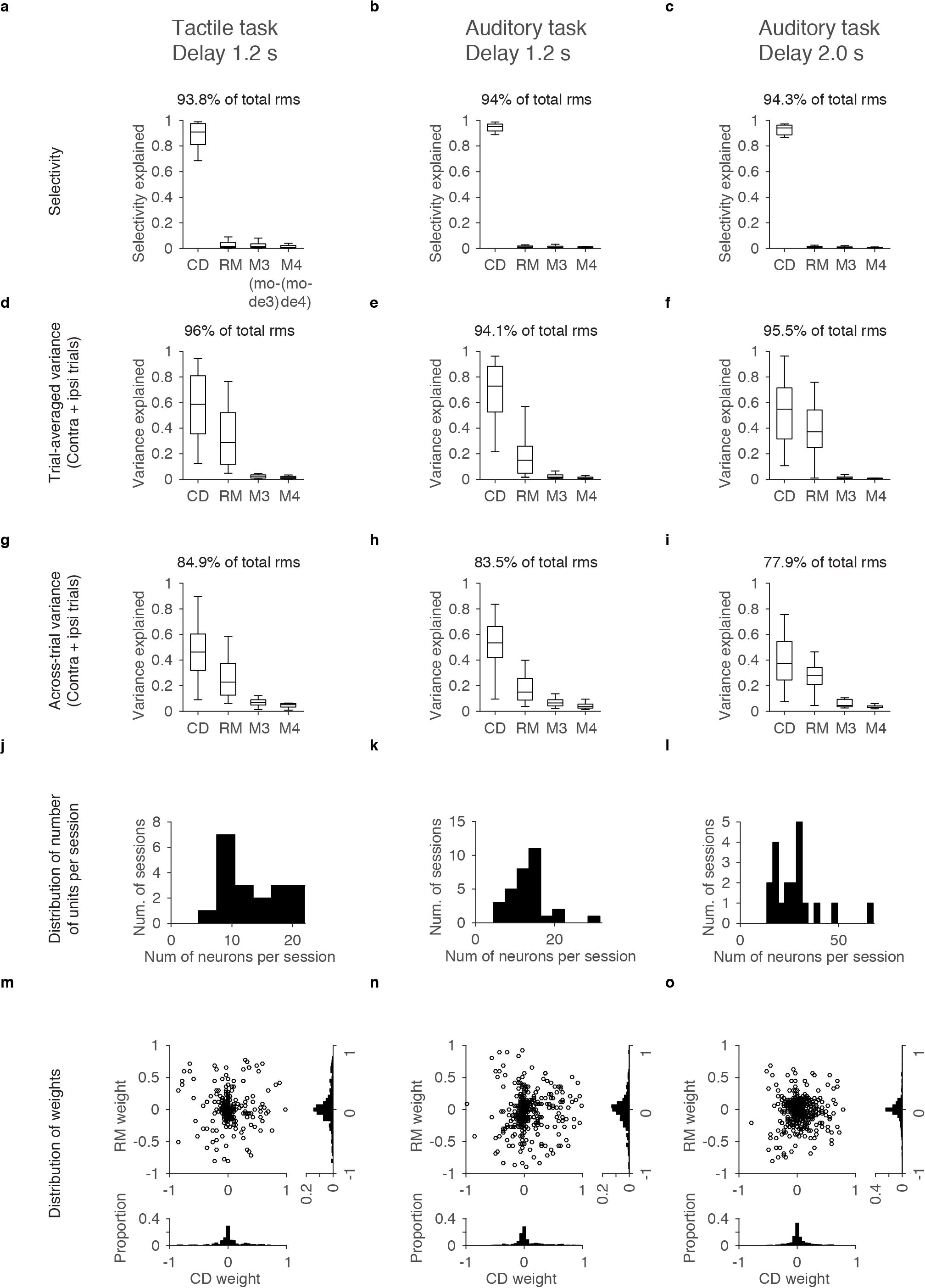
Variance explained by each mode. **a-c.** Selectivity explained by CD, RM, and remaining top two SVD modes (mode3, M3; mode 4, M4) (Methods). Sum of four modes are shown on top (mean across sessions). Central line in the box plot is the median. Top and bottom edges are the 75 % and 25 % points, respectively. The whiskers show the lowest datum within 1.5 interquartile range
(IQR) of the lower quartile, and the highest datum within 1.5 IQR of the upper quartile. **d-i** follow the same format. **d-f.** Trial-average variance explained by each mode (Methods). **g-i.** Across-trial variance explained by each mode (Methods). **j-l.** Distribution of number of putative pyramidal neurons simultaneously recorded. Because we used probes with less dead sites in the auditory task with 2.0 s delay, the yield was better. **m-o.** Distribution of CD and RM weights. Each point corresponds to single neuron. Neurons from all sessions were pooled. Right and bottom are histogram of RM weight and CD weight, respectively. Note that weights are widely distributed.

Fourth, we analyzed the remaining variance in activity in ALM. We performed singular value decomposition (SVD) of simultaneously recorded ALM neurons and orthogonalized the vectors to the CD. The projection of ALM activity to the first SVD component showed non-selective ramping (Fig. 15e-h) (‘ramping mode’, RM). CD and RM together explained more than 80 % of trial-averaged activity variance during the delay epoch (Fig. 16d-f, Methods). Over 60 % of across-trial variance was also explained by these two modes (Fig. 16g-i, Methods). The projections of ALM activity onto either CD or RM were ramping. This implies that, together with the variance measurements, the majority of activity in ALM can be explained as a weighted sum of selective ramping (CD) and non-selective ramping (RM). These analyses reveal that ALM activity is monotonic (ramping), and low dimensional (two dimensions capturing > 80 % of activity variance).

### Neurons with similar response show high spike count correlation

The low dimensionality of the across-trial variance implies that spike rates are correlated across neurons on a trial-by-trial level (spike count correlation, or noise correlation). Indeed, pairs of simultaneously recorded ramping-up cells showed similar activity patterns on a trial-by-trial basis (Fig. 17a). The onset and amplitude of ramping varied across trials within a cell, but was correlated among cells within a trial (Fig. 17a).

**Figure 17.**
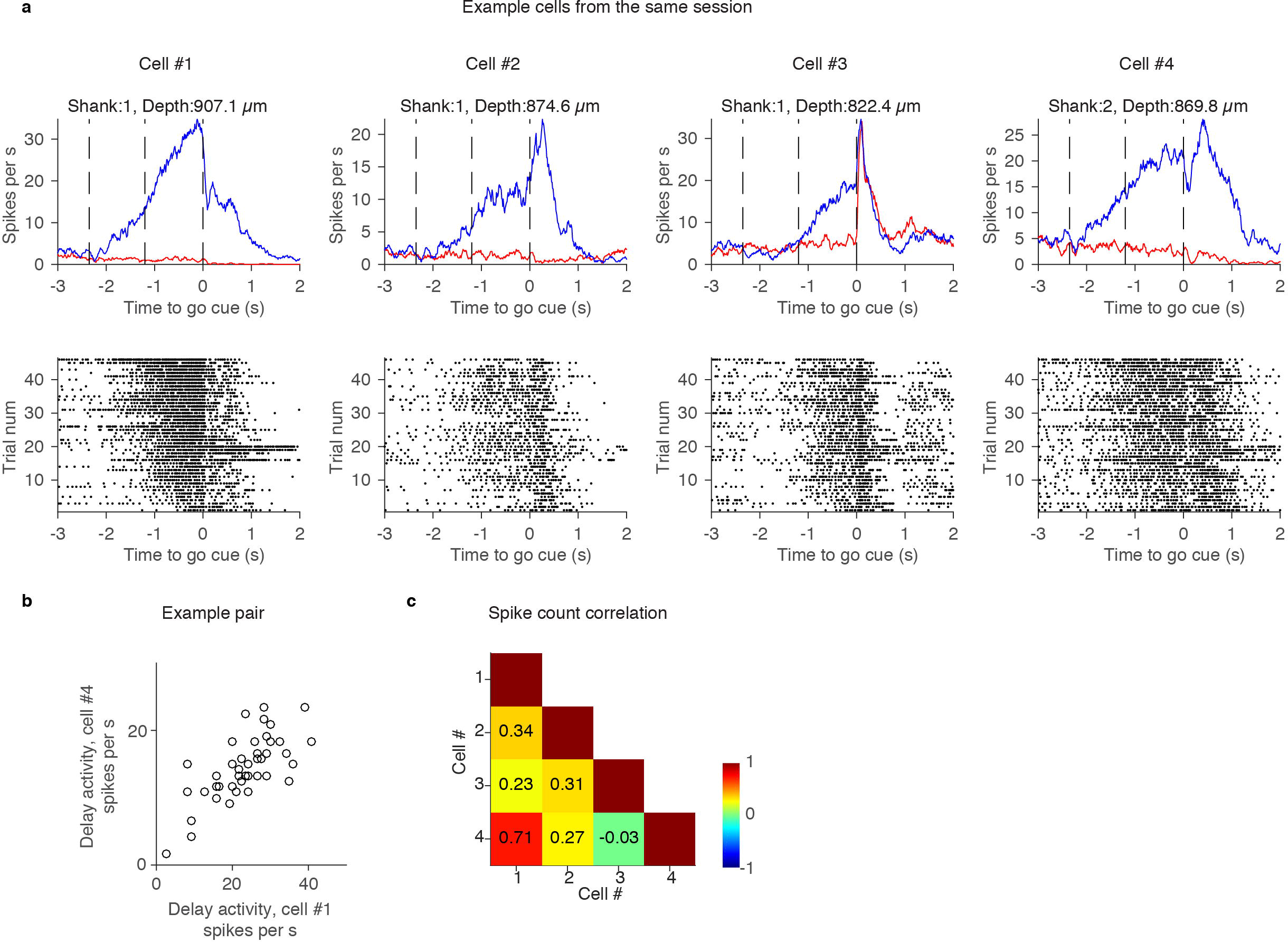
Example spike count correlation. **a.** Top, grand mean PSTH of ramping-up cells from the same session. Shank and the depth of each cell are shown; Bottom, spike raster in contra trials. Contra trials in all cells were sorted based on the rank order of spike count during the delay epoch in Cell #1. **b.** Relationship of contra delay activity (DA) in two example cells. Each circle represents a trial. **c.** Spike count correlation in contra trials calculated for all combinations of example cells. Grid color represents the value of spike count correlation (colorbar).

For each cell we defined a vector representing delay activity in each trial 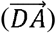. The length of 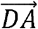 is the number of trials. To calculate spike count correlation, we measured the correlation between pairs of neurons as the Pearson’s correlation of 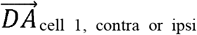 and 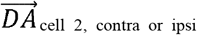. The example pairs of contra ramping-up cells showed high correlation of 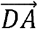 (Fig.17b, c).

We computed the correlation for all simultaneously recorded pairs. We classified cells into four major categories (following Fig. 7): contra ramping-up, ipsi ramping-up, ipsi ramping-down (or just “ramping-down” as contra ramping-down was rare) cells, and the others. Distributions of spike count correlations in contra trials between different types of cell pairs are shown (Fig. 18a-c, left column). The pairs between the same categories had positive correlation, whereas the pairs between different categories had correlation near 0 or slightly below 0. Spike count correlations in ipsi trials were similar (data not shown).

**Figure 18.**
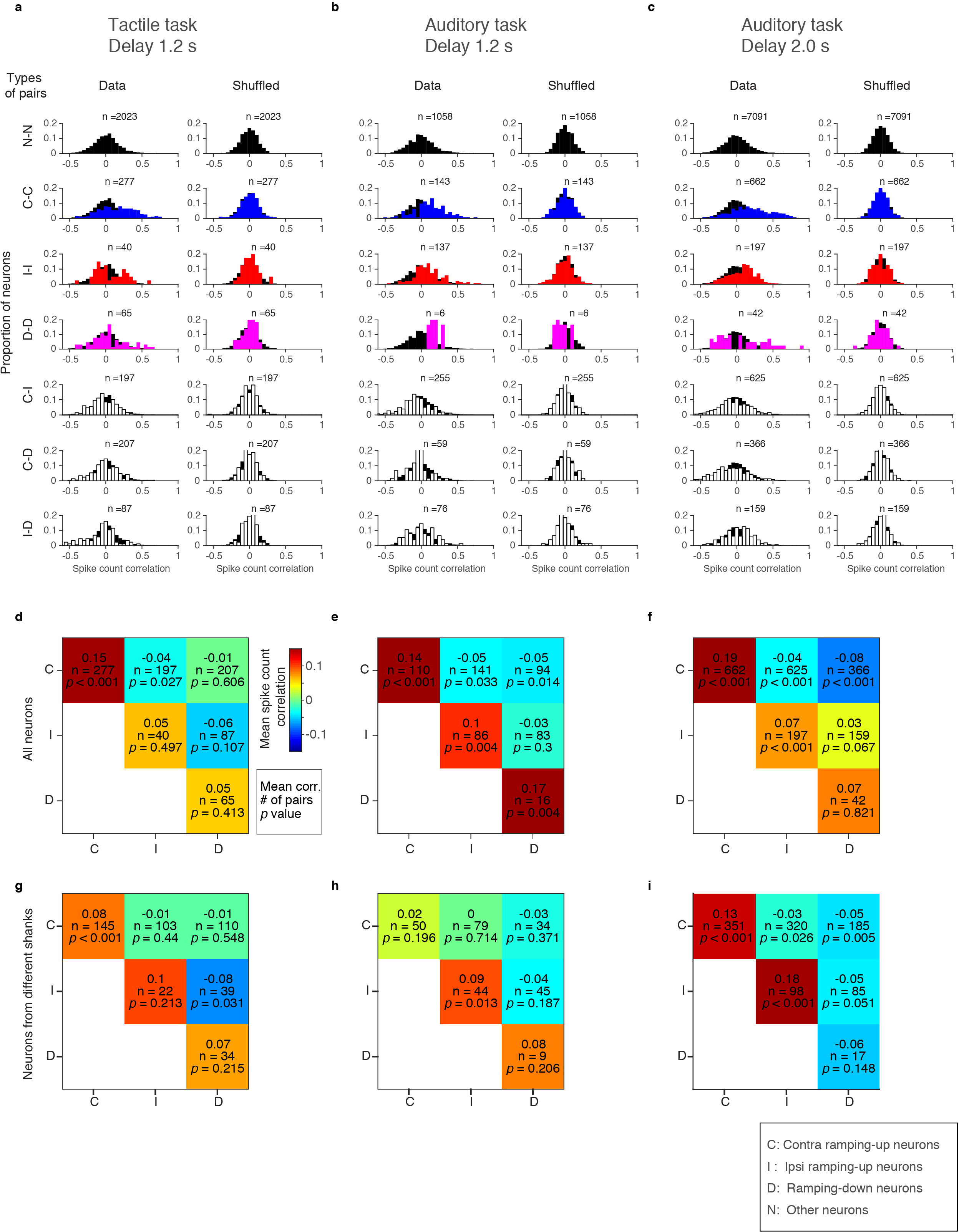
Spike count correlation. **a-c.** Distribution of spike count correlation between different subtypes of neurons in contra trials (left). Distribution is overlaid on top of the distribution of non-selective cells (black) for comparison. As a control, trials were shuffled for each neuron. Spike count correlation of the shuffled data is shown in the same format (right). C, contra-preferring ramping-up cells; I, ipsi-preferring ramping-up cells; D, ipsi-preferring ramping-down cells; N, other cells. Numbers of pairs are shown on top of the histogram. **d-f.** Mean spike count correlation between different subtypes of neurons in contra trials. Spike count correlation is shown in pseudo color (colormap, red: positive, blue: negative). Numbers of samples, and p-values (Methods) are shown below the noise correlation in each grid. **g-i.** Mean noise correlation between different subtypes of neurons from different shanks.

As a control, we shuffled trial labels within the same trial type and calculated the spike count correlation (Fig. 18a-c, right column). This procedure destroyed across-trial correlation among neurons without changing the mean PSTH of individual neurons. Comparison of the data with shuffled controls (Fig. 18a-c, right column) provides a probability for the measured correlation to be higher than chance level (*p*-values in Fig. 18d-f, *Mann–Whitney U* test).

This measure confirms that spike count correlations are significantly high between neurons within categories. Correlations across categories are in some cases negative (Fig. 18d-f). The results were similar even when we computed the correlation for pairs from different shanks (shank interval: 250 μm) (Fig. 18g-i). High spike count correlation among cells with the same coding preference is consistent with data from frontal eye fields of primates during motor-planning ^33^.

## Discussion

We compared three different delayed-response tasks to identify common features of neural activity underlying motor planning. In all cases ALM was necessary for motor planning (Fig. 2). Although there was heterogeneity in the dynamics across individual ALM neurons (Fig. 4), most features of the dynamics during the delay epoch were similar across tasks and similar to findings in non-human primates and rats. First, the majority of ALM neurons showed significant selectivity during the delay epoch (Fig. 5, 6) ^3-5^. Second, intermingled neurons were selective for either contra or ipsi movements (Fig. 5, 6) ^3–5,9^. Third, the ramping dynamics of contra-selective and ipsi-selective neurons were markedly different (Fig. 7, 8) ^4,9^. The proportion of ramping-up cells and absolute value of delay activity were higher for contra-selective neurons than ipsi-preferring neurons. Fourth, many cells showed mixed coding, with a bias toward preparatory activity (Fig. 10, 11) ^8,9,28^. Fifth, the amplitude of ramping activity was similar, independent of the delay durations (Fig. 6) ^4,45^, suggesting that the rate of ramping is determined by the timing of the action ^22,46^. Sixth, ALM activity was monotonic and low-dimensional (Fig.13-16) ^31^. Seventh, spike count correlations were high among cells with the same coding preference (Fig. 17-18) ^33^. These findings suggest that features of motor planning are common across tasks with different instruction cues and actions and also across multiple species. Moreover, similar neural circuit mechanisms likely mediate motor planning independent of sensory modalities and actions.

Recent advances in neurophysiological recording methods (high-density silicon probe recordings and calcium imaging) enable us to monitor tens to hundreds of neurons simultaneously. Such population-recordings provide information not available from single cell recoding ^47^. First, since neural activity (and calcium response caused by spikes) is highly variable, trial averaging is often necessary to infer the coding of neurons, often preventing analysis of single trials. Large scale recordings allow population averaging and provide the statistical power to analyze circuit activity at the level of single trials. This is critical to understand the relationship between neuronal activity and behavior. For example, population analysis in premotor cortex allows prediction of movement direction ^30^ and action timing ^16,30^. Second, information encoded in the correlation structure of network activity is not interpretable by monitoring single neurons. Dimensionality reduction methods allow us to extract the “modes” coding for the variable of interest, such as licking direction ^30^. Based on dimensionality reduction, we found that over 80 % of activity during the delay epoch was explained by two modes. The low dimensionality may be due to simplicity of the task ^31^. Testing how the dimensionality changes in tasks with more behavioral choices is an important goal for the future.

Sequentially activated neurons during motor planning and other kinds of short-term memory tasks have been reported in premotor cortex, mPFC, PPC and hippocampus of rodents ^10,42-44^. These observations provide support for a theoretical model explaining persistent activity with feedforward or recurrent networks that effectively extend network time constants ^41,48-53^. On the other hand, monotonic ramping observed in our data excludes such model to explain preparatory activity in ALM. The lack of sequential activity might be due to difference in task or brain region.

Multiple pyramidal cell types have been reported in ALM based on gene expression profile and anatomical properties, such as long-range axonal projection ^54^. These diverse defined cell types are likely to show different activity patterns due to distinct anatomical connections and intrinsic properties. For example, pyramidal tract neurons that project out of cortex show stronger contra laterality compared to the intratelencephalic neurons that project to other cortical areas ^29^. Cell type specific recordings to test how the features of delay activity differ among cell types is another important goal for the future.

High spike count correlations among neurons sharing the same coding preference indicated strong functional connection between them, or sharing of common inputs. The strong functional connection can be mediated by local recurrent connections. Alternatively it can be mediated by strong long-range excitatory loop between ALM and thalamus (thalamo-cortical loop) ^55^. Such functional network structures alone is not sufficient to reveal the possible mechanisms underlying motor-planning, as the same network structure can result for instance in an integrator or attractor networks ^56^. To further nail down the mechanism of motor planning, optogenetic perturbations are necessary to identify the circuit mechanisms of preparatory activity. Motor planning in mice, which is similar to that in non-human primates, enable us to interrogate the circuits in order to unveil the underlying mechanism.

## Acknowledgements

We thank Nuo Li, Tim Wang, Liu Liu and Arseny Finkelstein for comments on the manuscript, Tim Harris, Brian Barbarits, Jun Jaeyoon James and Wei-Lung Sun for help with silicon probe recordings and spike sorting, and Lorenzo Fontolan and Shaul Druckmann for discussions. This work was funded by Howard Hughes Medical Institute. H.K.I is a Helen Hay Whitney Foundation postdoctoral fellow.

## Author Contributions

H.K.I. performed experiments and analyzed data, with input from K.S.. M.I. supported animal trainings and performed inactivation screen. R.S. provided theoretical feedbacks. H.K.I. and K.S. wrote the paper, with input from all the authors.

## Author Information

The authors declare no competing interests. Correspondence and requests for materials should be addressed to.

## Methods

### Mice

This study is based on 27 mice (age > P60, male). We used four transgenic mouse lines: PV-IRES-cre ^37^, Olig3-cre ^57^, Ai32 (Rosa-CAG-LSL-ChR2(H134R)-EYFP-WPRE, JAX 012569) ^38^, VGAT-ChR2-EYFP ^58^. Eight PV-ires-cre ×Ai32 mice were used for behavioral experiments (Fig.2). Nineteen mice were used for recordings (Supplementary file). Some of these mice were in addition used for various optogenetic experiments not described in this paper {Inagaki et al, *in prep*}. Because optogenetic manipulations in one trial did not affect performance and spike rate in the next trial (Table 2 and 3), we pooled data from these different experiments after excluding trials with optogenetic stimulation.

All procedures were in accordance with protocols approved by the Janelia Institutional Animal Care and Use Committee. Detailed information on water restriction, surgical procedures and behavior have been published ^35^. All surgical procedures were carried out aseptically under 1-2 % isofluorane anesthesia. Buprenorphine HCl (0.1 mg/kg, intraperitoneal injection; Bedford Laboratories) was used for postoperative analgesia. Ketoprofen (5 mg/kg, subcutaneous injection; Fort Dodge Animal Health) was used at the time of surgery and postoperatively to reduce inflammation. After the surgery, mice were allowed free access to water for at least three days before start of water restriction. Mice were housed in a 12:12 reverse light:dark cycle and behaviorally tested during the dark phase. A typical behavioral session lasted 1 to 2 hours and mice obtained all of their water in the behavior apparatus (approximately 1 ml per day; 0.3 ml was supplemented if mice drank less than 0.5 ml). On other days mice received 1 ml water per day. Mice were implanted with a titanium headpost. For ALM photoinhibition, mice were implanted with a clear-skull cap ^8^. Craniotomies for recordings were made after behavioral trainings.

### Behavior

For the tactile task, at the beginning of each trial, a metal pole (diameter, 0.9 mm) moved within reach of the whiskers (0.2 s travel time) for 1.0 second, after which it was retracted (0.2 s retraction time) (Fig. 1a). The sample epoch (1.4 s total) was the time from onset of the pole movement to completion of the pole retraction (Fig. 1a). The delay epoch lasted for another 1.2 seconds after completion of pole retraction. An auditory ‘go’ cue separated the delay and the response epochs (pure tone, 3.4 kHz, 0.1 s).

For the auditory task (Fig. 1b), at the beginning of each trial, five repetitive tones were presented in one of two frequencies: 3 or 12 kHz. Each tone was played for 150 ms with 100 ms between tones. The sample epoch (1.15 s total) was the time from onset of the first tone to completion of the last tone (Fig. 1b). The delay epoch lasted for another 1.2 or 2.0 s after completion of the last tone. An auditory ‘go’ cue with intermediate frequency (carrier frequency 6kHz, with 360 Hz modulating frequency to make it distinct from instruction tones) separated the delay and the response epochs (0.1 s). To compensate the sound intensity for tuning curve of C57BL6 auditory system^59^, the sound pressure was 80, 70, and 60 dB for 3, 6 and 12 kHz sound, respectively. These frequencies are relatively invulnerable to hearing loss observed in C57BL6 mice^60^.

In both tasks, a two-spout lickport (4.5 mm between spouts) was used to record licking events and deliver water rewards. After the ‘go’ cue, licking the correct lickport produced a water reward (approximately 2 μL); licking the incorrect lickport triggered a timeout (0-5 s). Licking early during the trial (‘lick early’ trials) triggered a timeout (1 s). Trials in which mice did not lick within 1.5 s after the ‘go’ cue (‘no response’ trials) were rare and typically occurred at the end of behavioral sessions. ‘No response’ trials and ‘lick early’ trials were excluded from analyses. For trainings, we started with short delay (0 s for the tactile task, 0.3 s for the auditory task) and gradually increased the delay duration (Fig. 1c).

### Photoinhibition

Photoinhibition was deployed on 33-50 % (Fig. 2b) or 25 % (Fig. 2cd) of behavioral trials. To prevent mice from distinguishing photoinhibition trials from control trials using visual cues, a ‘masking flash’ (40 × 1 ms pulses at 10 Hz) was delivered using 470 nm LEDs (Luxeon Star) near the eyes of the mice throughout the trial. Photostimuli from a 473 nm laser (Laser Quantum) were controlled by an acousto-optical modulator (AOM; Quanta Tech) and a shutter (Vincent Associates).

Photoinhibition of ALM was performed through a clear-skull cap (beam diameter at the skull: 400 μm at 4 σ) ^8^. The light transmission through the intact skull is 50 % ^8^. We stimulated parvalbumin-positive interneurons expressing ChR2 in PV-IRES-Cre mice crossed to Ai32 reporter mice (Fig. 2). Behavioral and electrophysiological experiments showed that photoinihibition in the PV-IRES-Cre × Ai32 mice was indistinguishable from the VGAT-ChR2-EYFP mice ^55^. Photoinhibition silences a cortical area of 1 mm radius (at half-max) through all cortical layers ^8^.

To silence cortex unilaterally during the sample or delay epoch (Fig. 2b, c), we photostimulated for 1.2 s, including the 200 ms ramp, starting at the beginning of the sample or delay epoch. We used 40 Hz photostimulation with a sinusoidal temporal profile (1.5mW average power) and a 200 ms linear ramp during laser offset (this reduced rebound neuronal activity). We used scanning Galvo mirrors to inactivate different locations of the cortex. For the screen (Fig. 2a), photoinhibition location was randomly selected for each trial. For Fig. 2c, we photoinhibited ALM (anterior 2.5 mm, lateral 1.5 mm, bregma) or S1 (posterior 1.5 mm, lateral 3.5 mm, bregma).

To silence ALM bilaterally (Fig. 2d), we photostimulatd four spots centering ALM or M1 (anterior 0 mm, lateral 1.5 mm, bregma) in each hemisphere with 1mm spacing (in total 8 spots) using scanning Galvo mirrors. We photoinhibited each spot sequentially, with 5ms per step. We photostimulated with constant laser power with the 200 ms ramp at the end. Mean laser power was 1.5 mW per spot.

### Behavioral data analysis

Behavioral performance was the fraction of correct trials, excluding ‘lick early’ and ‘no response’ trials. Early lick rate was the fraction of lick early trials, excluding ‘no response’ trials.

For statistics (Fig. 2), we performed hierarchical bootstrapping: first we randomly selected animals with replacement, second randomly selected sessions within an animal with replacement, and last randomly selected trials within the session with replacement ^8,61-63^. We counted in how many bootstrap trials, the trials with particular photostimulation have lower performance than control trials. If less than 2.5 % or more than 97.5 % trials were higher, we concluded that the stimulation resulted in significant effect (*α* = 0.05).

### Extracellular electrophysiology

A small craniotomy (diameter, 0.5 mm) was made over the left ALM hemisphere one day prior to the recording session. Extracellular spikes were recorded using Janelia silicon probes with two shanks (250 μm between shanks) (Part# A2×32-8mm-25-250-165). The 64 channel voltage signals were multiplexed, recorded on a PCI6133 board (National instrument) and digitized at 400 KHz (14 bit). The signals were demultiplexed into the 64 voltage traces sampled at 25 kHz and stored for offline analysis. One to five recording sessions were obtained per craniotomy. Recording depth was inferred from manipulator readings (Table 1). The craniotomy was filled with cortex buffer (125 mM NaCl, 5 mM KCl, 10 mM glucose, 10 mM HEPES, 2 mM MgSO4, 2 mM CaCl2. Adjust pH to 7.4) and the brain was not covered. The tissue was allowed to settle for at least five minutes before the recording started.

### Extracellular recording data analysis

The extracellular recording traces were band-pass filtered (300-6k Hz). Events that exceeded an amplitude threshold (4 standard deviations above the background) were sorted using JRclust ^64^.

Units were classified based on spike width. Spike widths were computed as the trough-to-peak interval in the mean spike waveform. Units with width < 0.35 ms were defined as putative fast-spiking GABAergic interneurons (FS neurons), and units with width > 0.5 ms as putative pyramidal neurons. Units with intermediate spike widths (0.35 – 0.5 ms) were excluded from our analyses. This scheme was verified by optogenetic tagging of parvalbumin-positive neurons (data not shown) ^8^. See Table 1 and supplementary file for number of sessions and units. Only putative pyramidal neurons were analyzed for all the figures, except Fig. 9.

Neurons were tested for trial type selectivity during the sample, delay, and response epochs by comparing spike counts during contra-and ipsi-trials in each epoch (*Mann–Whitney U* test, *p* < 0. 05; Fig. 5). To compute “contra-selectivity” for each neuron, we calculated spike rate difference between the contra and ipsi trials (*SR*_*t,contra*_ – *SR*_*t,ipsi*_, where *SR* denotes spike rate at each time point *t* in each trial type) (Fig. 3a). To compute “normalized contra-selectivity” we normalized the contra selectivity by the peak value for each neuron (Fig. 5a-c).

Neurons with selectivity during the delay epoch were further classified into “contra-preferring” versus “ipsi-preferring”, based on their mean spike rate during the delay epoch (Fig. 3a and 6). Selectivity is the absolute spike rate difference between the two trial types (see Fig. 3a for detail, Fig. 6d-f). We defined “delay activity (DA)” as difference of spike rate between the baseline presample epoch versus delay epoch in each neuron (*SR*_*delay,contra or ipsi*_ – *SR*_*pre-sample,contra or ipsi*_) (Fig. 3a, 7-9). We referred to neurons with increasing delay activity in preferred direction as ramping-up, and decreasing delay activity in preferred direction as ramping-down cells (*Mann–Whitney U* test, *P* < 0.05; Fig. 3a and 7).

For the peri-stimulus time histograms, only correct trials were included, unless specified (Fig. 11e-h). Spikes were averaged over 100ms with 1ms step. Bootstrapping was used to estimate standard errors of mean.

### Angle analysis *(Fig. 10, 11*)

To test what cells are coding (sensory input and/or movement), we compared selectivity in the correct trials and incorrect trials. We defined the *r* and *θ* as below:

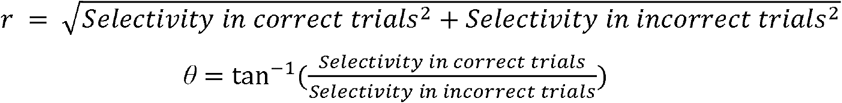

We analyzed neurons with more than 10 incorrect trials for each trial-type (incorrect lick-right (IR) and incorrect lick-left (IL) trials). Cells with *r* < 2 were excluded from the analysis of *θ*. The standard error of mean (*bootstrap*) of the angle measurement were 17.4⍰ and 18.3⍰ for the tactile task, and auditory task, respectively.

### ROC analysis *(Fig. 10g, h*)

For ROC analysis, we analyzed neurons with more than 10 incorrect trials for each trial-type (IR and IL trials). For each cell, we randomly subsampled 10 trials each of correct lick right (CR) trials, correct lick left (CL) trials, IR trials, and IL trials. To decode sensory input, we pooled CR and IR trials together, and CL and IL together. We computed ROC curve for each neuron and calculated the area under the curve (Fig. 10g, h). To decode motor output, we pooled CR and IL trials together, and CL and IR together. Area under the curve lower than 0.5 was subtracted from 1. Chance level performance was calculated by shuffling trial types (CR, CL, IR and IL). This permutation has been repeated 1000 times for each cell to determine 95 % confidence interval.

### Toy models *(Fig. 12*)

We modeled 50 neurons for each scenario. For the scenario in which each neuron shows ramping activity (Fig. 12a), the spike rate of each neuron linearly ramps from 0 to range between 3/5 ~ 30 spikes per s (the peak spike rate is evenly distributed). For the scenario in which each neuron shows bump-like activity (Fig. 12b), each neuron follows the activity pattern (*f*_*n*_) described below:

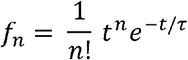

Here, **τ** denotes time constant of each neuron (25ms). This is linear filtering of a pulse input for n times (The same equation used in Fig. 1 in the Goldman, 2009 ^41^). In both cases, we added Gaussian noise (σ = 3 spikes per s), and averaged over 10 ms (Fig. 12c, d). We calculated rank correlation, and stability of CD (Fig. 12e-h) in the same way as we processed the data (see below).

### Peak analysis *(Fig. 13*)

For each neuron with delay selectivity, we randomly subselected half of the contra trials to calculate mean spike rate (as in other analysis spike rates were averaged over 100 ms). We identified the time point with max spike rate during the delay epoch. Similarly we calculated mean spike rate for the remaining trials and identified the peak timing. We repeated this procedure 1000 times. Density of peak location between two halves of trials was shown in pseudo color (Fig. 13g-i). As a control, we shuffled timing of spikes in each trial within the delay epoch. After shuffling we followed the same procedures described above. This procedure was repeated for 1000 times. Results of analyzing ipsis trial were similar (data not shown). Results of analyzing all pyramidal neurons were similar, too (data not shown).

### Population vector analysis *(Fig. 15-16*)

For the session based analysis (Fig. 15 and 16), recording sessions with more than 5 simultaneously recorded pyramidal cells were analyzed (Supplementary file). To calculate coding direction (CD) for a population of n simultaneously recorded neurons, we found an n × 1 vector maximally distinguishing two trial types (contra and ispi trials), in the n dimensional activity space. To compare CD over time (Fig. 15i-k), we calculated average spike rates in contra and ipsi trials separately for each neuron. Then, 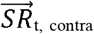 and 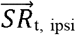 are n × 1 vectors of average spike rates at a time point *t* (10 ms step). The difference in the mean spike rate vector, 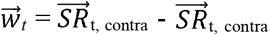, was divided by its own norm. This is CD at the time point. Pearson’s correlations between CD across time points were shown in Fig. 15i-k.

Since correlation of CD was high during the delay epoch, for other figures (Fig. 15a-c and Fig. 16) we averaged 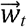 during the last 600 ms of the delay epoch and normalized by its own norm to obtain the CD. We calculated CD based on randomly selected half of trials. To obtain trajectories along CD, we projected the spike rate in the remaining trials to the CD as an inner product (Fig. 15a-c). To calculate the “selectivity explained” (Fig. 16a-c), we first calculated the total selectivity as a square sum of the selectivity across neurons (square sum of n × 1 vector). Then we calculated the square of selectivity of the projection along the CD, and divided it by the total selectivity. Here, selectivity was calculated based on the last 600 ms of the delay epoch. To calculate the “trial-average variance” (Fig. 16d-f), we first calculated the total variance as a square sum of the mean delay activity (the last 600 ms of the delay epoch) across neurons in each trial type (contra and ipsi trial) (square sum of n × 2 matrix). Then we calculated the square sum of the mean delay activity in projection along the CD. We divided this value by the total variance. To calculate “across-trial variance” (Fig. 16g-i), we first calculated the total variance as a square sum of the delay activity (the last 600 ms of the delay epoch) across neurons and across trials (square sum of n × trial-number matrix). Then we calculated the square sum of the delay activity in projection along the CD across trials. We divided this value by the total variance.

To find modes explaining the remaining activity variance (RM, M3 andM4), we first found eigenvectors of the population activity matrix using singular value decomposition (SVD) at each time point and averaged over the last 600 ms of the delay epoch. The data for SVD at each time point was n × trial-number matrix. We analyzed the first three eigenvectors from the SVD. All of these vectors were rotated using the Gram-Schmidt process to be orthogonal to CD and to each other. Since the projection to the first vector resulted in non-selective ramping activity (Fig. 15f-h), we referred to this vector as a ramping mode. The other two vectors are named mode 3 (M3) and mode 4 (M4).

### Spike count correlation *(Fig. 17-18*)

To calculate spike count correlation among neurons, we first calculated delay activity vector 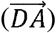 of each neuron in contra trials. 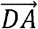 is trial-number × 1 vector, consisting of delay activity in each trial. Then we calculated Pearson’s correlation between 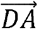 vectors of different neurons recorded in the same session. Neurons were classified into four classes: Contra ramping-up neurons, ipsi ramping-up neurons, ramping-down neurons (Contra ramping-down neurons were excluded from analysis because of small number, Fig. 7) and all the other neurons. As a control, we shuffled trial label in each neuron independently to remove across cell correlation without affecting mean spike rate of each neuron. To calculate *p*-value, we performed *Mann–Whitney U* test (two-sided) comparing spike-count correlations of the data and the shuffled data.

### Statistics and data

The sample sizes are similar to sample sizes used in the field (more than few hundreds units per task). No statistical methods were used to determine sample size. We did not exclude any animal for data analysis. Trial types were randomly determined by a computer program during the experiment. During spike sorting, experimenters cannot tell the trial type, so experimenters were blind to conditions. All bootstrapping was done over 1,000 iterations, unless otherwise described. Data sets will be shared at CRCNS.ORG in the NWB format.

